# Trajectories for the evolution of bacterial CO_2_-concentrating mechanisms

**DOI:** 10.1101/2022.06.21.497102

**Authors:** Avi I. Flamholz, Eli Dugan, Justin Panich, John J. Desmarais, Luke M. Oltrogge, Woodward W. Fischer, Steven W. Singer, David F. Savage

**Affiliations:** Department of Molecular and Cell Biology, University of California, Berkeley, California 94720, United States; Division of Biology and Biological Engineering, California Institute of Technology, Pasadena, CA 91125; Resnick Sustainability Institute, California Institute of Technology, Pasadena, CA 91125, USA; Biological Systems and Engineering Division, Lawrence Berkeley National Laboratory, Berkeley, CA 94720, USA; Division of Geological & Planetary Sciences, California Institute of Technology, Pasadena, CA 91125; Howard Hughes Medical Institute, University of California, Berkeley, California 94720

**Keywords:** physiology, evolution, photosynthesis, Earth history, synthetic biology

## Abstract

Cyanobacteria rely on CO_2_ concentrating mechanisms (CCMs) that depend on ≈15 genes to produce two protein complexes: an inorganic carbon (Ci) transporter and a 100+ nm carboxysome compartment that encapsulates rubisco with a carbonic anhydrase (CA) enzyme. Mutations disrupting CCM components prohibit growth in today’s atmosphere (0.04% CO_2_), indicating that CCMs evolved to cope with declining environmental CO_2_. Indeed, geochemical data and models indicate that atmospheric CO_2_ has been generally decreasing from high concentrations over the last ≈3.5 billion years. We used a synthetic reconstitution of a bacterial CCM in *E. coli* to study the co-evolution of CCMs with atmospheric CO_2_. We constructed strains expressing putative ancestors of modern CCMs — strains lacking one or more CCM components — and evaluated their growth in a variety of CO_2_ concentrations. Partial forms expressing CA or Ci uptake genes grew better than controls in intermediate CO_2_ levels (≈1%); we observed similar phenotypes in genetic studies of two autotrophic bacteria, *H. neapolitanus* and *C. necator.* To explain how partial CCMs improve growth, we advance a model of co-limitation of autotrophic growth by CO_2_ and HCO_3_^-^, as both are required to produce biomass. Our model and results delineated a viable trajectory for bacterial CCM evolution where decreasing atmospheric CO_2_ induces an HCO_3_^-^ deficiency that is alleviated by acquisition of CAs or Ci uptake genes, thereby enabling the emergence of a modern CCM. This work underscores the importance of considering physiology and environmental context when studying the evolution of biological complexity.

**Significance:** The greenhouse gas content of the ancient atmosphere is estimated using models and measurements of geochemical proxies. Some inferred high ancient CO_2_ levels using models of biological CO_2_ fixation to interpret the C isotopes found in preserved organic matter. Others argued that elevated CO_2_ could reconcile a faint young Sun with evidence for liquid water on Earth. We took a complementary “synthetic biological” approach to understanding the composition of the ancient atmosphere by studying present-day bacteria engineered to resemble ancient autotrophs. By showing that it is simpler to rationalize the emergence of modern bacterial autotrophs if CO_2_ was once high, these investigations provided independent evidence for the view that CO_2_ concentrations were significantly elevated in the atmosphere of early Earth.

## Introduction

Nearly all carbon enters the biosphere through CO_2_ fixation in the Calvin-Benson-Bassham (CBB) cycle. Rubisco is the carboxylating enzyme of that pathway and the most abundant enzyme on Earth (1, 2). Rubisco is often considered inefficient due to relatively slow carboxylation kinetics (3–5) and non-specific oxygenation of its five-carbon substrate ribulose 1,5-bisphosphate, or RuBP (6, 7). However, rubisco arose more than 2.5 billion years ago, when the Earth’s atmosphere contained virtually no O_2_ and, many argue, far more CO_2_ than today (8, 9). Over geologic timescales, photosynthetic O_2_ production (8) and CO_2_-consuming silicate weathering reactions (10) are thought to have caused a gradual increase in atmospheric levels of O_2_ (≈20% of 1 bar atmosphere today) and depletion of atmospheric CO_2_ to present-day levels of a few hundred parts per million (≈280 ppm pre industrial, ≈420 ppm or ≈0.04% today). The amount of atmospheric CO_2_ during the Archean Eon (4-2.5 Ga) is challenging to estimate accurately from the geological record, but is thought to have been substantially higher than today, perhaps as high as 0.1-1 bar (9, 11, 12). It is likely, therefore, that contemporary autotrophs grow on drastically lower levels of CO_2_ than their ancestors did.

Many photosynthetic organisms evolved CO_2_ Concentrating Mechanisms (CCMs), which help meet the challenge of fixing carbon in a low CO_2_ atmosphere. CCMs concentrate CO_2_ near rubisco and are found in all Cyanobacteria, some Proteobacteria, as well as many eukaryotic algae and diverse plants (13). Because CO_2_ and O_2_ addition occur at the same active site in rubisco (6), elevated CO_2_ has the dual effects of accelerating carboxylation and suppressing oxygenation of RuBP by competitive inhibition (14–16). As shown in Figure 1A, bacterial CCMs are encoded by ≈15 genes comprising three primary features: (i) an energy-coupled inorganic carbon (Ci) transporter at the cell membrane and (ii) a cytosolic 100+ nm protein compartment called the carboxysome found that (iii) co-encapsulates rubisco with a carbonic anhydrase (CA) enzyme (4, 17). Energized Ci transport produces a high HCO_3_^-^ concentration in the cytosol (≈30 mM, Fig. 1A), which is converted into a high carboxysomal CO_2_ concentration by carbonic anhydrase activity, localized exclusively to the carboxysome (16, 18, 19).

**Fig. 1:**
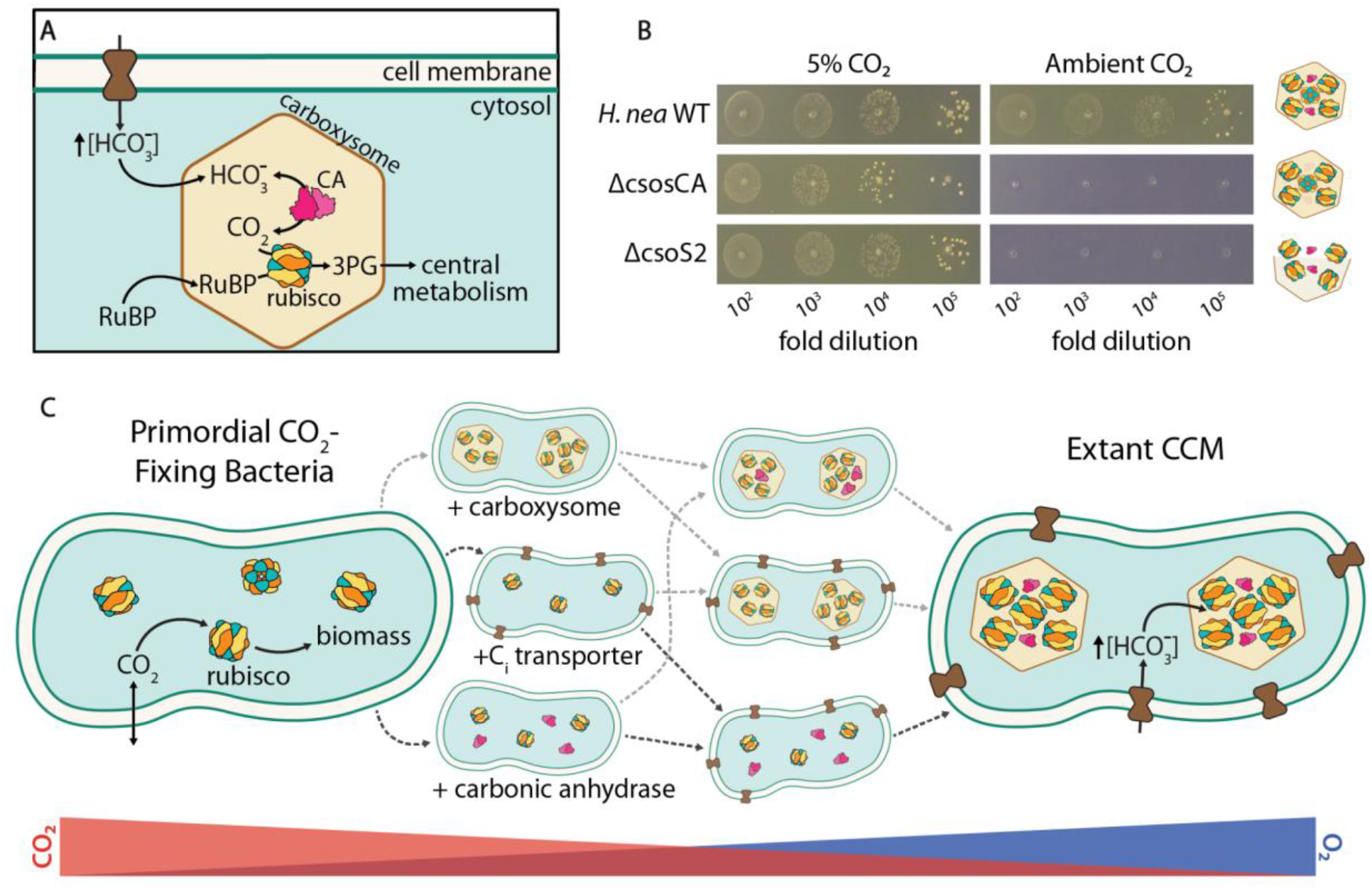
Mechanism and potential routes for the evolution of the bacterial CO_2_ concentrating mechanism. (A) Today the bacterial CCM functions through the concerted action of two protein complexes-an inorganic carbon (Ci) transporter at the cell membrane and carboxysome encapsulation of rubisco with carbonic anhydrase (CA). Ci uptake leads to a high intracellular HCO_3_^-^ concentration, well above equilibrium with the external environment. Elevated HCO_3_^-^ is converted to a high carboxysomal CO_2_ concentration by CA activity located only there, which promotes carboxylation by rubisco. (B) Mutants lacking genes coding for essential CCM components grow in elevated CO_2_ but fail to grow in ambient air, exemplified here by the chemoautotroph *H. neapolitanus.* Strains lacking the carboxysomal CA (Δ*csosCA*) or an unstructured protein required for carboxysome formation (Δ*csos2*) failed to grow in ambient air, but grew robustly in 5% CO_2_ (> 10^8^ colony forming units/ml, Figure S1). See Table S4 for description of mutant strains. (C) We consider the CCM to be composed of three functionalities beyond rubisco itself: a CA enzyme (magenta), an Ci transporter (dark brown) and carboxysome encapsulation of rubisco (light brown). Since atmospheric O_2_ levels were extremely low (≈1 ppm, blue triangle) and CO_2_ was likely quite high (perhaps 0.1-1 bar, red triangle) during the Archean epoch, primordial CO_2_-fixing bacteria would not have needed a CCM. We sought to discriminate experimentally between the six sequential trajectories (dashed arrows) in which CCM components could have been acquired.

CCM genes are straightforward to identify as mutations disrupting essential CCM components prohibit growth in ambient air (Fig. 1B) and mutants are typically grown in 1% CO_2_ or more (14, 15, 20–22). At first glance, therefore, the CCM appears to be “irreducibly complex” as all plausible recent ancestors - e.g. strains lacking individual CCM genes - are not viable in the present-day atmosphere. Irreducible complexity is incompatible with evolution by natural selection, so we and others supposed that bacterial CCMs evolved over a protracted interval of Earth history when atmospheric CO_2_ concentrations were much greater than today (13, 23–26). We therefore hypothesized that ancestral forms of the bacterial CCM (i.e. those lacking some genes and complexes required today) would have improved organismal fitness in the elevated CO_2_ environments that prevailed when they arose.

To test the hypothesis, we reconstructed present-day analogues of plausible CCM ancestors (henceforth “analogues of ancestral CCMs”) and tested their growth across a range of CO_2_ partial pressures. Our goal was to identify a stepwise pathway of gene acquisition supporting the evolutionary emergence of a bacterial CCM by improving growth in ever-decreasing CO_2_ concentrations (Fig. 1C). We focused on trajectories involving sequential acquisition of genetic components because carbonic anhydrases (27), Ci transporters (22), and homologs of carboxysome shell genes (28, 29) are widespread among bacteria and could therefore be acquired horizontally.

One approach to constructing contemporary analogues of CCM ancestors is to remove CCM genes from a native host. If CCM components were acquired sequentially, some single gene knockouts would be analogous to recent ancestors, e.g. those lacking a complete carboxysome shell (30). We tested this approach by assaying a whole-genome knockout library of a γ-proteobacterial chemoautotroph, *H. neapolitanus,* in five CO_2_ partial pressures (22, 31). As shown in Figure 2 and elaborated below, we found that many CCM genes contribute substantially to organismal fitness even at CO_2_ concentrations tenfold greater than present-day atmosphere (0.5% CO_2_), supporting the view that CCM components play an important physiological role even in relatively high environmental CO_2_ concentrations.

**Fig. 2:**
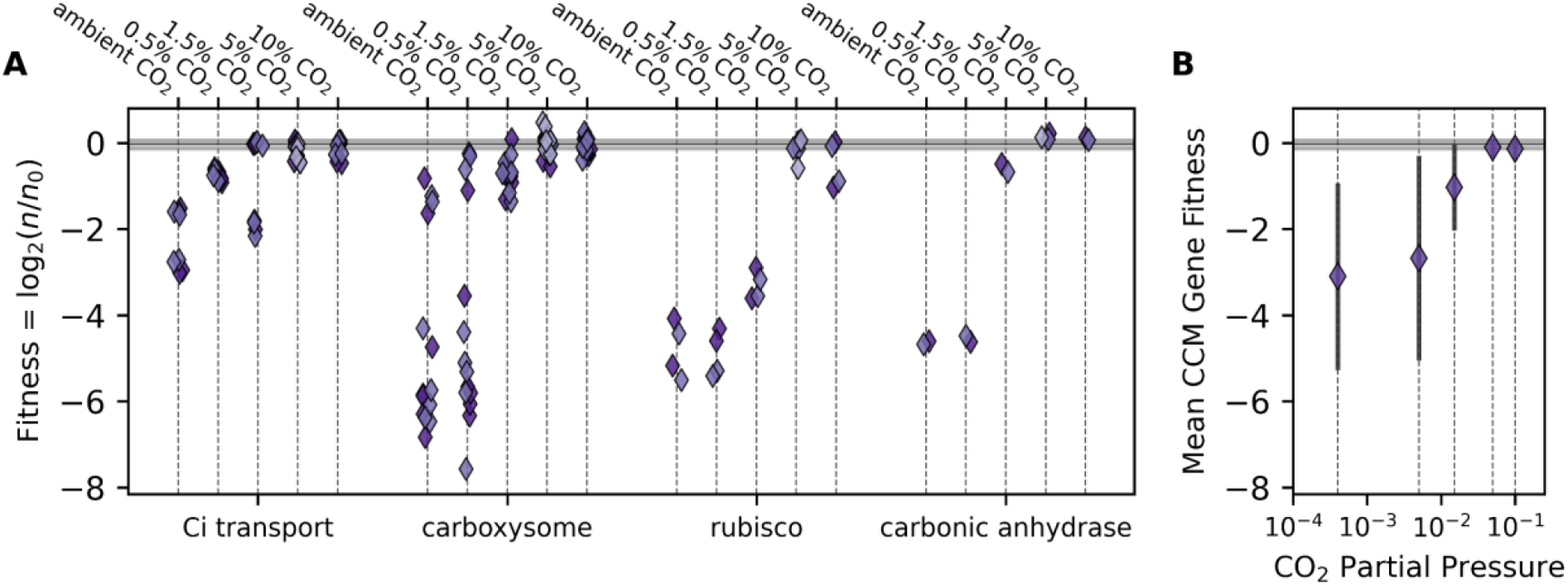
The contribution of *H. neapolitanus* CCM genes to organismal fitness depends on the environmental CO_2_ concentration. *H. neapolitanus* is a chemoautotroph that natively utilizes a CCM in low CO_2_ environments. (A) Using a barcoded transposon library constructed in this organism (22), we profiled the contributions of CCM genes to organismal fitness during autotrophic growth across a range of CO_2_ levels *(Methods).* Fitness is estimated as the log_2_ ratio of barcode counts between the endpoint sample and the pre-culture for each mutant (22, 32). Each point gives the average fitness of multiple mutants to a single CCM gene in a given CO_2_ condition. The library contained 30 mutants disrupting the carboxysomal CA gene (*csosCA*) and an average of ≈40 mutants per CCM gene. A fitness of - 2 implies that gene disruption was, on average, associated with a fourfold decrease in strain abundance after a defined period of growth. Biological replicates are indicated by shading, and the gray bar gives the interquartile range of fitness effects for all ≈1700 mutants across all CO_2_ levels (−0.15−0.065). “Ci transport” includes 4 DAB genes in two operons (22), “carboxysome” includes 6 nonenzymatic carboxysome genes, “rubisco” denotes the two subunits of the carboxysomal rubisco, and “CA” the carboxysomal CA gene. *H. neapolitanus* also expresses a secondary rubisco, which explains why disruption of the carboxysomal rubisco is not lethal in high CO_2_ (Figure S3). (B) On average, CCM genes contribute less to organismal fitness as environmental CO_2_ levels rise. Error bars give the standard deviation of fitness values across all CCM genes. See Figures S2-3 for analysis of reproducibility and Tables S1-3 for detailed description of genes.

Removing single CCM genes from a native host can only produce analogues of recent ancestors, however. We recently constructed a functional CCM in an engineered *E. coli* strain called CCMB1 (4). This strain depends on rubisco carboxylation for growth and the expression of a full complement of CCM genes enabled growth in ambient air. Here we used CCMB1 to construct analogues of ancestral CCMs, including several lacking two or more essential components of modern CCMs. We assayed the growth of these putative ancestors across a range of CO_2_ pressures to determine whether any ancestral forms contribute to organismal fitness — i.e. improve growth relative to a control strain expressing only rubisco — in various CO_2_ pressures. In the following sections we describe the growth of these strains and discuss how it can inform our entwined understandings of bacterial physiology, CCM evolution and the CO_2_ content of Earth’s ancient atmosphere.

## Results and Discussion

### CCM genes contribute to fitness even in elevated CO_2_

Using a barcoded genome-wide transposon mutagenesis screen, we previously demonstrated that a 20-gene cluster in *H. neapolitanus* contains all the genes necessary for a functional CCM (4, 22). The original screen measured gene fitness across the entire genome via batch competition assays, comparing the abundance of strains in high CO_2_ (5%) and ambient air (0.04%) via high-throughput sequencing (22). This effort demonstrated that at least 12/20 genes are necessary components of the CCM as disruption of any one produced substantial and reproducible growth defects in ambient air. To mimic the changes in atmospheric CO_2_ that likely occurred over Earth history, we quantified the same library in three additional CO_2_ pressures to cover five CO_2_ levels: ambient (≈0.04% CO_2_), low (0.5%), moderate (1.5%), high (5%), and very high (10%). Replicate experiments were strongly correlated (R > 0.85, Figure S2), implying a high degree of reproducibility. We therefore proceeded to ask whether *H. neapolitanus* CCM genes contribute to fitness in elevated CO_2_ conditions.

Figure 2 plots the effect of disrupting CCM genes across five CO_2_ pressures, with genes grouped by their known roles in the CCM. The library contained an average of ≈35 mutants per gene in the *H. neapolitanus* genome (22), so each point in Figure 2A represents the average fitness of 5-50 individual mutants. Surprisingly, we found that many CCM genes also contributed substantially to organismal fitness in 0.5% and 1.5% CO_2_, indicated by large fitness defects (negative values in Figure 2A), resulting in a negative average fitness of CCM mutants when the CO_2_ pressure was 1.5% or less (Figure 2B). Carboxysome genes, for example, were critical for growth in 0.5% CO_2_, while certain Ci transport genes contributed substantially to growth in 0.5% and 1.5% CO_2_.

Though several *H. neapolitanus* CCM genes contributed to fitness in intermediate CO_2_ levels, the CCM appeared to be dispensable in high CO_2_ (5-10%, Fig. 2B). As such, the data presented in Figure 2 indicated that individual CCM components like the carboxysome, CA, or Ci transporter may improve autotrophic growth in intermediate CO_2_ levels (≈1%) even in the absence of a complete and functional CCM. We considered testing this hypothesis in *H. neapolitanus* directly by constructing “ancestral-like” CCMs lacking carboxysome shell genes or Ci transporters. However, genetic manipulation of *H. neapolitanus* is cumbersome, and the native host’s regulatory network could complicate interpretation — a concern highlighted by three DNA-binding proteins that emerged in our screen as likely regulators of the CCM (Figure S3 and Table S1). We therefore decided to construct and test analogues of CCM ancestors in a non-native host, namely *E. coli*.

### Evaluating putative ancestral CCMs in a rubisco-dependent *E. coli*

We recently developed an *E. coli* strain, CCMB1, that depends on rubisco carboxylation for growth in minimal medium. This strain requires elevated CO_2_ for rubisco-dependent growth, but expressing the *H. neapolitanus* CCM from two plasmids enabled growth in ambient air (4). One of these plasmids, pCB’, encodes the carboxysome genes along with the encapsulated rubisco and carbonic anhydrase enzymes (4). The other plasmid, pCCM’, encodes the DAB1 inorganic carbon transporter (4, 22), two rubisco chaperones (22, 33, 34) and a carboxysome positioning system (35). We used this two-plasmid system to express analogues of putative CCM ancestors in CCMB1 and assayed these strains’ growth in ambient air, 0.5%, 1.5% and 5% CO_2_. We compared the growth of strains expressing partial CCMs to that of a reference CCMB1 strain expressing the *H. neapolitanus* rubisco on the p1A vector but no other CCM components (4). The reference strain grew robustly in 5% CO_2_ but failed to grow in ambient air (Fig. 3A, ‘Rubisco Alone’).

**Fig. 3:**
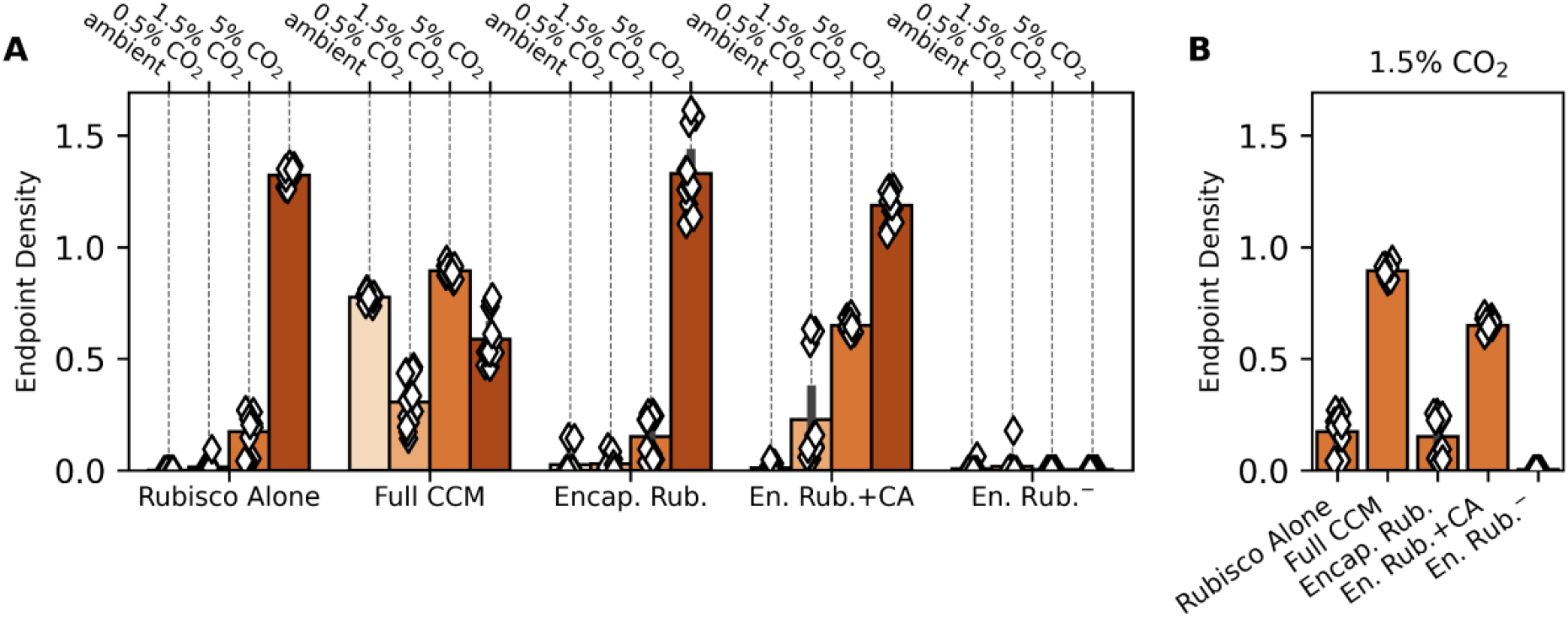
Encapsulation of rubisco alone does not improve rubisco dependent growth of CCMB1 *E. coli* in any CO_2_ level tested. (A) Using our reconstitution of the *H. neapolitanus* CCM in *E. coli* (4) we tested whether rubisco encapsulation has any effect on growth at four CO_2_ partial pressures (Methods). Each diamond represents one technical replicate of four biological replicates. The CCMB1 strain grows in elevated CO_2_ (1.5 and 5%) when rubisco is expressed (“Rubisco Alone”, left). As previously reported, expressing the full complement of CCM genes from the pCB’ and pCCM’ plasmids (“Full CCM”) enabled growth in all CO_2_ levels. By replacing pCCM with a vector control and making an inactivating mutation to the carboxysomal CA (CsosCA C173S) we were able to express rubisco in a carboxysome without CA or Ci transport activities (“Encap. Rub.”). This strain grew similarly to the reference “Rubisco Alone” in all conditions. When the CA active site was left intact (“En. Rub.+CA”), growth improved above the “Rubisco Alone” baseline in 0.5% and 1.5% CO_2_. Finally, a negative control strain carrying inactive rubisco (“En. Rub.^-^”, CbbL K194M) failed to grow in all CO_2_ conditions. (B) Focusing on the growth in 1.5% CO_2_ highlights the contribution of CA activity to rubisco-dependent growth. See Tables S4-5 for description of strains and plasmids, see Figures S4-5 for growth curves and analysis of statistical significance.

Consistent with our previous work (4), expression of the full complement of CCM genes permitted growth in all CO_2_ concentrations tested (Fig. 3A, ‘Full CCM’). Recent modeling efforts endeavored to support the hypothesis that encapsulation of rubisco in a semi-permeable barrier could improve CO_2_ fixation by generating an acidic local pH (26). This proposal does not require Ci transport or CA activity and, if correct, would therefore support an “encapsulation first” model of CCM evolution. To evaluate the effect of encapsulating rubisco alone in a protein compartment, we replaced the pCCM’ plasmid with a vector control so that no Ci transporter was expressed and further deactivated the carboxysomal CA by mutating a crucial cysteine residue (pCB’ CsosCA C173S, ‘Encap. Rub.’ in Fig. 3A). Encapsulating rubisco on its own did not improve growth over the reference strain in any CO_2_ concentration tested. However, when we left the CA active site intact (‘En. Rub.+CA’), growth improved substantially in intermediate CO_2_ levels (0.5% and 1.5%). These data indicated that CA plays a pivotal role in autotrophic growth in low CO_2_, but did not support the hypothesis that encapsulation alone improves CO_2_ fixation by rubisco (26).

### Carbonic anhydrase and energy-coupled inorganic carbon transport improve the growth of rubisco-dependent E. coli

Our observation that carboxysomal CA activity improves rubisco-dependent growth on its own (i.e. without also expressing a Ci transporter) motivated us to test the effects of CA and Ci transport independently of carboxysome expression. We designed plasmids that express the *H. neapolitanus* Dab2 Ci transporter (22), *E. coli’s* native CA (Can) or both. This was achieved by cloning both Dab2 and Can into a dual expression vector and making active site mutants (DabA C539A, Can C48A, D50A) to isolate each activity. These vectors were co-transformed into CCMB1 with a constitutive version of p1A (p1Ac) so that rubisco expression would not be affected by induction of Dab2 or Can *(Methods).*

Expressing active Can and Dab2, whether alone or together, improved growth substantially in 1.5% CO_2_ (compare ‘+DAB+CA^-^’, ‘+DAB^-^+CA’, and ‘+DAB+CA’ to ‘Rubisco Alone’ in Figure 4). This effect was even more pronounced in 0.5% CO_2_, where the reference strain failed to grow (Figures 4A and S7). Similar to the reference strain, a double-negative control strain expressing inactivated versions of both Dab2 and Can (‘+Dab^-^+CA^-^’) grew poorly or not at all in 0.5% and 1.5% CO_2_, implying that the observed growth improvements were due to activity and not a side-effect of heterologous gene expression.

**Fig. 4:**
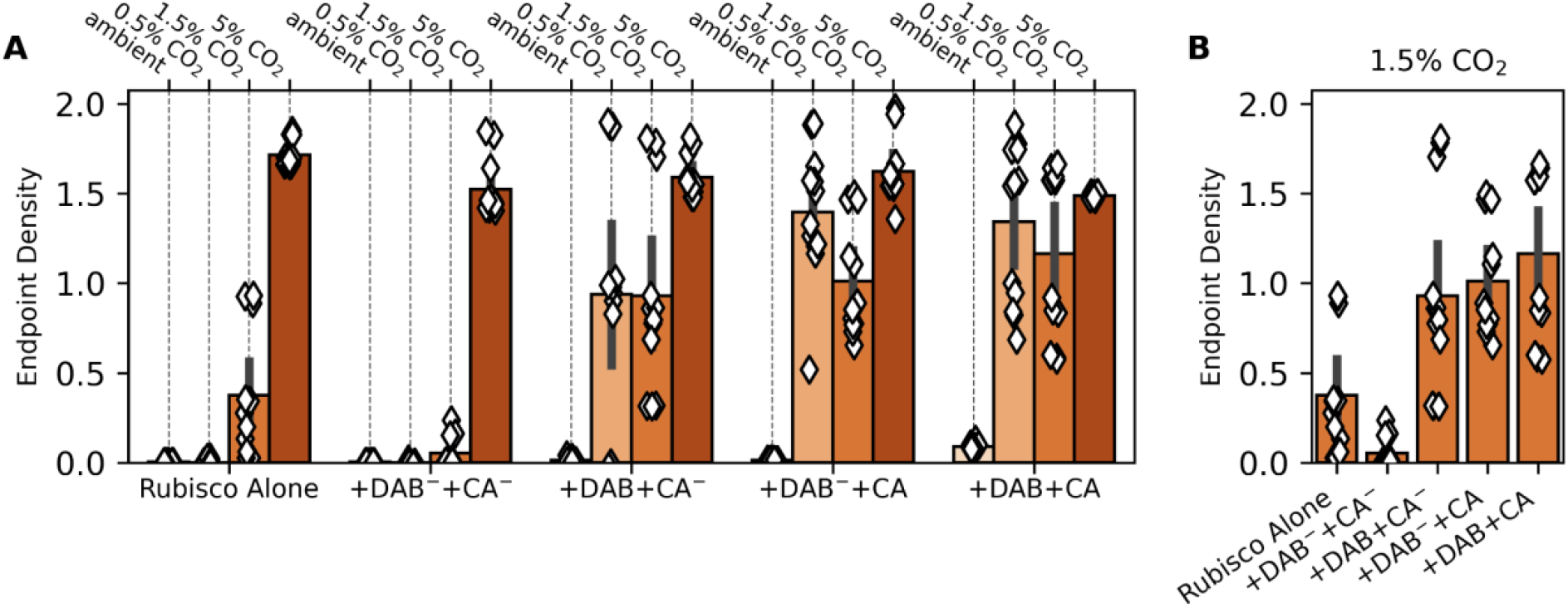
Expression of carbonic anhydrase or Ci transport improves rubisco-dependent growth of CCMB1 *E. coli* in intermediate CO_2_ levels even in the absence of other CCM components. As in Figure 3, the reference strain (“Rubisco Alone”) grew only in elevated CO_2_ (1.5% and 5%). The remaining strains expressed the *E. coli* carbonic anhydrase (Can) and the DAB2 Ci transporter from a second plasmid. These activities were isolated by means of active site mutations. The negative control “+DAB^-^ +CA^-^” expressed inactive Can (C48A, D50A) and DAB2 (DabA2 C539A) and grew less robustly than the reference in all conditions. If either active site was left intact (“+DAB+CA^-^” or “+DAB^-^+CA”) we observed a sizable growth improvement in both 0.5 and 1.5% CO_2_. This growth improvement remained when both active sites were left intact (“+DAB+CA”). Panel (B) emphasizes this effect clearly by focusing on growth in 1.5% CO_2_. See Table S4 for strain genotypes, Figures S6-7 for growth curves and analysis of statistical significance.

It was not immediately obvious to us how CA and Ci uptake activities improve rubisco-dependent growth of CCMB1 *E. coli.* We found the effects of Dab2 expression especially perplexing because Ci uptake is expected to generate intracellular HCO_3_^-^ (16–18, 22, 36) while the rubisco substrate is CO_2_ (37). To confirm that these results are not a side-effect of working in an engineered *E. coli* strain, we pursued genetic experiments in a natively autotrophic proteobacterium, *C. necator.*

### *C. necator* depends on CA for autotrophic growth at intermediate pCO_2_

While all photosynthetic Cyanobacteria rely on the CBB cycle and a full complement of CCM genes (38), some chemoautotrophic bacteria depend on the CBB cycle but lack identifiable genes encoding carboxysome components or Ci transporters (39–42). As most characterized bacterial rubiscos are not CO_2_-saturated in ambient air and are, in addition, substantially inhibited by atmospheric levels of O_2_ (7), we expected that such organisms would require elevated CO_2_ for robust growth.

*Cupriavidus necator,* formerly known as *Ralstonia eutropha,* is one such bacterium (43, 44). *C. necator* is a facultative chemolithoautotroph typically found at the interface between oxic and anoxic environments where H_2_ and O_2_ coexist. Such “knallgas” environments include soils, sediments and geothermal sites (45) that are often characterized by elevated CO_2_ (45–47). While *C. necator* is an obligate aerobe capable of chemoautotrophic growth on H_2_, CO_2_ and O_2_ via the CBB cycle, it has no carboxysome genes and no obvious Ci transporters (41). Consistent with previous reports (48), autotrophic growth of wild-type *C. necator* was very poor in ambient CO_2_ (‘*C. necator* WT’ in Figure 5A). We generated a double CA knockout strain, *C. necator Δcan Δcaa* (42, 49) and found that CA removal greatly attenuated autotrophic growth in 0.5% and 1.5% CO_2_ (‘*C. necator* ΔCA’). Consistent with our experiments in *E. coli,* this growth defect was complemented by heterologous expression of a human CA (‘ΔCA+CA’) or a DAB-type Ci uptake system (‘ΔCA+DAB’). Moreover, as in *E. coli,* co-expression of Ci uptake with native CAs was not deleterious (‘WT+DAB’). Rather, this strain grew to higher densities than wild-type in 1.5% and 5% CO_2_ (e.g. Fig. 5B).

**Fig. 5:**
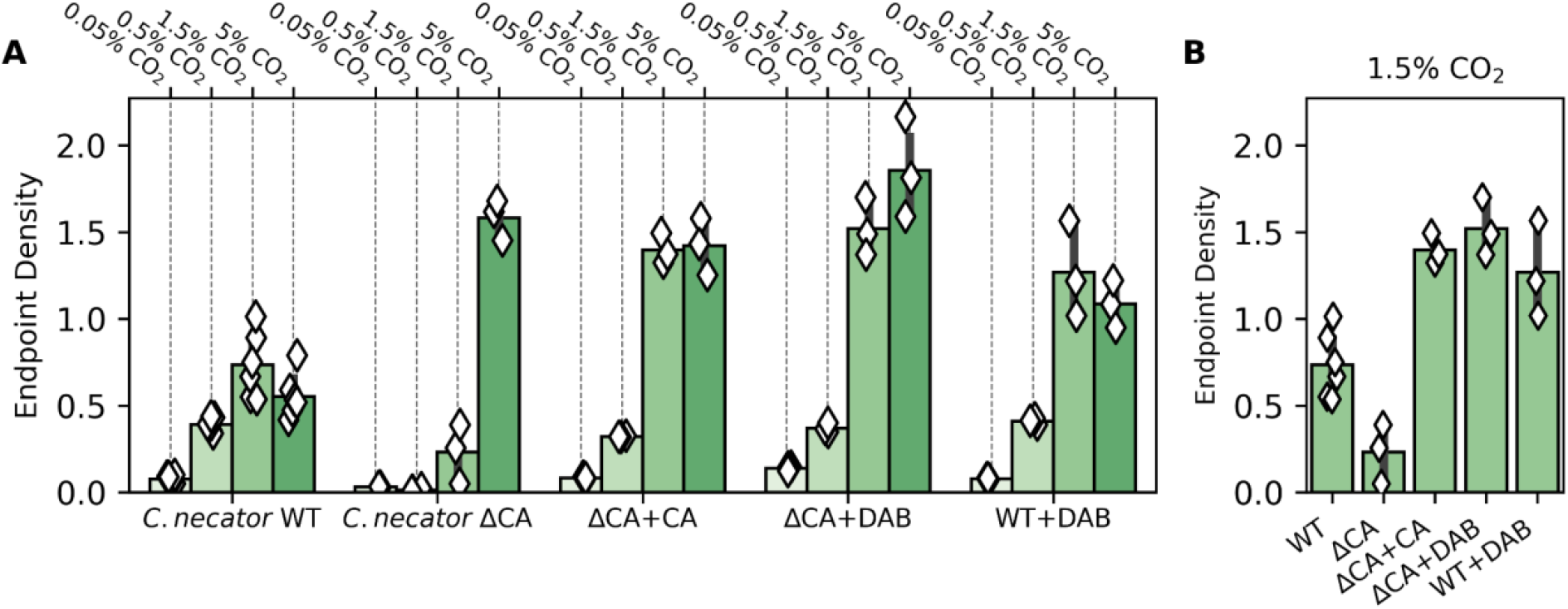
*C. necator* requires carbonic anhydrase or Ci uptake for robust autotrophic growth in 0.5% and 1.5% CO_2_. *C. necator* strains were grown autotrophically in minimal medium at a variety of CO_2_ levels and endpoint optical density was measured after 48 hours *(Methods).* (A) Growth of the *C. necator* double carbonic anhydrase knockout (ΔCA) was greatly impaired growth in 0.5% and 1.5% CO_2_. Panel (B) focuses on 1.5% CO_2_. Compared to wild-type *C. necator* (WT), which grew to a final OD600 of 0.73±0.28 in 1.5% CO_2_ (6 biological replicates), growth of ΔCA was greatly impaired, reaching a final OD of 0.23±0.17 (3 biological replicates). Expression of either the human carbonic anhydrase II (ΔCA+CA) or the DAB2 Ci transporter from *H. neapolitanus* (ΔCA+DAB) recovered robust growth which exceeded even the wild type, indicating that the wild type may not express saturating levels of CA. See Figure S8 for statistical analysis.

### A nutritional requirement for HCO_3_^-^ explains the observed phenotypes

We found that expression of a CA, Ci transporter, or both improved rubisco-dependent growth of *C. necator* and CCMB1 *E. coli* in intermediate CO_2_ concentrations (0.5 and 1.5%, Figs. 4–5). Furthermore, to our surprise, a CCMB1 strain expressing active Dab2 and Can grew reproducibly, if slowly, in ambient air (‘+Dab+CA’ in Figure 6). This was remarkable because biological membranes are very permeable to CO_2_, with a permeability coefficient P_c_ ≈ 0.1-1 cm/s (16, 50, 51) that is several orders-of-magnitude larger than those of small charged species like acetate, phosphate and bicarbonate (50–53). Given this high permeability, we expected co-expression of energized HCO_3_^-^ uptake with a CA to generate a deleterious “futile cycle” where energy expended pumping HCO_3_^-^ was wasted when CO_2_ produced by CA activity “leaks” back out of the cell. Instead we found that such a cycle is compatible with rubisco-dependent growth of two bacteria in relatively low CO_2_ (1.5% or lower).

**Fig. 6:**
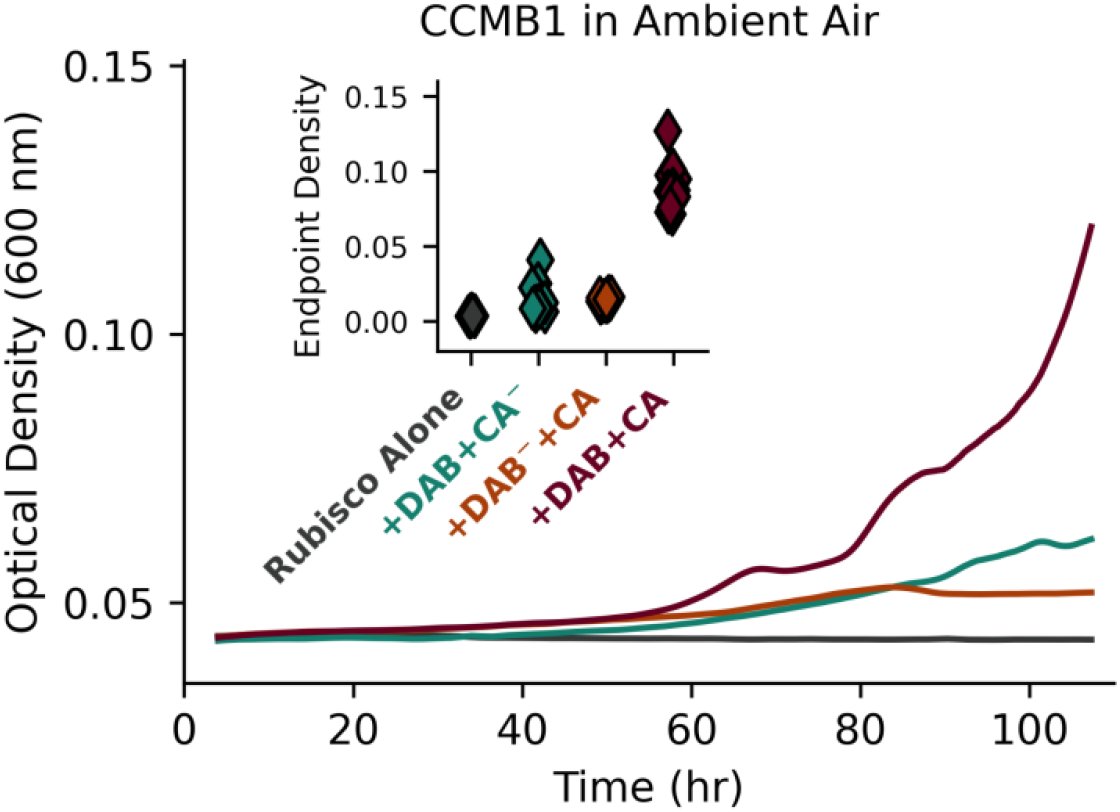
Co-expression of carbonic anhydrase and Ci uptake enabled rubisco-dependent growth of CCMB1 *E. coli* in ambient air. Inspecting the ambient CO_2_ growth data presented in Figure 4 revealed that coexpression of CA and Ci transport (‘Rub.+DAB+CA’) substantially improved rubisco-dependent growth of CCMB1 *E. coli* in ambient CO_2_ concentrations. This effect was modest (≈0.1 OD units above the ‘Rubisco Alone’ control) but reproducible, as indicated by endpoint data plotted on the inset. Curves are colored to match labels on the inset. See Figure S9 for statistical analysis.

A naive explanation of how CA expression might improve growth is that it increases the intracellular CO_2_ concentration relative to the reference strain expressing only rubisco. For this hypothesis to explain the benefit of CA expression, rubisco activity must deplete intracellular CO_2_ (C_in_) significantly below its extracellular level (C_out_) in the reference strain. Otherwise CA can have no effect, as carbonic anhydrases are not coupled to any energy source and so cannot cause C_in_ to exceed C_out_. However, this naive model cannot explain the growth benefits associated with expressing Ci uptake systems, which provide HCO_3_^-^ and not CO_2_. Moreover, the following calculation shows that this hypothesis is unreasonable precisely because CO_2_ is so membrane-permeable that rubisco cannot deplete C_in_ much beneath C_out_.

In a bacterium, rubisco might make up 20% of soluble protein at the very most. This amounts to a mass concentration of roughly 0.2 x 300 mg/ml ≈ 60 mg/ml rubisco (54). As each rubisco active site is attached to ≈60 kDa of protein (BNID <u>105007</u>), the maximum rubisco active site concentration is ≈1 mM. In this naive model, C_in_ is set by the balance of passive uptake through the membrane, with effective permeability *a* = P_c_ x SA/V ≈ 10^4^ s^-1^, and fixation by rubisco, with an effective rate constant of *γ* = [rubisco] x k_cat_/K_M_ < 10^3^ s^-1^. These values give a steady-state C_in_ of *α* C_out_ / (*α + γ*) > 0.9 C_out_. That is, even an extreme level of rubisco activity cannot draw C_in_ beneath 90% of C_out_. In such conditions, rubisco would fix ≈10^10^ CO_2_/hour, supporting a 1-2 hr doubling time (see *Supplementary Information* for full calculation). So, although the CO_2_ concentration is low in aqueous environments equilibrated with present-day atmosphere (≈10 μM), passive CO_2_ uptake can support substantial rubisco flux and, therefore, CO_2_ limitation of rubisco carboxylation cannot explain the observed phenotypes.

We argue that the effects of expressing CA or Ci transport can be explained by the ubiquitous dependence of growth on HCO_3_^-^ (55–57). It has been clear for at least 80 years that heterotrophs also require inorganic carbon for growth. Seminal investigations in the 1930s and 1940s advanced the hypothesis that this dependence is due to a specific requirement for HCO_3_^-^ (58–61), which is the substrate of several carboxylation reactions involved in lipid, nucleotide and arginine biosynthesis. This explanation is now supported by detailed experiments in several heterotrophs demonstrating that growth in ambient air can be supported either by CA activity (55–57) or by providing the products of central-metabolic carboxylation reactions in the growth media (56, 57). The metabolic networks of chemoautotrophs, Cyanobacteria and plants imply an equivalent dependence on HCO_3_^-^ for biosynthesis (62–64). We therefore advance a model of co-limitation of autotrophic growth by both CO_2_ and HCO_3_^-^-dependent carboxylation fluxes, where most of biomass carbon derives from rubisco-catalyzed carboxylation of CO_2_ and a small minority derives from HCO_3_^-^ (Figure 7A). We formalized this notion as an idealized mathematical model fully described in *Supplementary Information*.

**Fig. 7:**
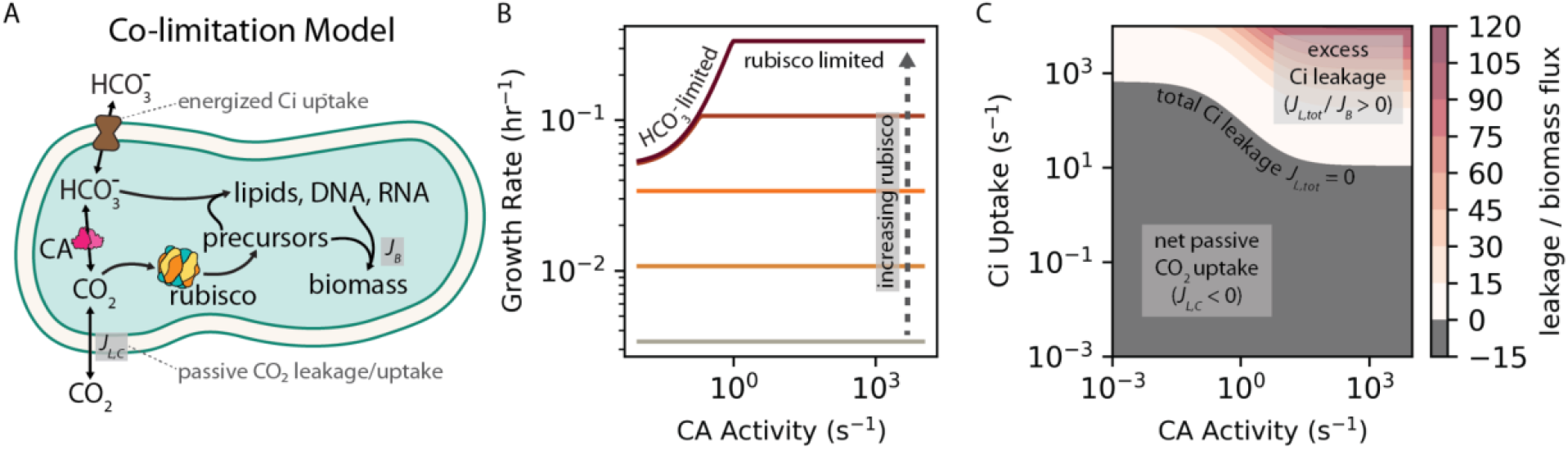
Co-limitation of autotrophic growth by CO_2_ and HCO_3_^-^-dependent carboxylation reactions explains the growth improvements associated with expressing carbonic anhydrases and Ci transporters. (A) In autotrophs using the CBB cycle, nearly all biomass carbon derives from rubisco-catalyzed CO_2_ fixation. However, autotrophs also require HCO_3_^-^ for carboxylation reactions in lipid, nucleic acid, and arginine biosynthesis (55–57). We expressed this diagram as a mathematical model, which we applied to understand why CA and Ci uptake improved rubisco-dependent growth. (B) The model exhibited two regimes: one wherein growth was limited by rubisco flux another where it was limited by HCO_3_^-^-dependent carboxylation (“bicarboxylation”) flux. At low rubisco levels (lighter-colored lines), growth was rubisco-limited: increased rubisco activity produced faster growth, but the growth rate was insensitive to CA activity because slow CO_2_ hydration provided sufficient HCO_3_^-^ to keep pace with rubisco. At higher rubisco levels (maroon lines), growth was bicarboxylation-limited and increased CA activity was required for increasing rubisco activity to translate into faster growth. Increasing Ci uptake led to similar effects (Figure S11). In panel (C), color indicates the ratio of total Ci leakage (*J_L,tot_* = *J_L,C_* + *J_L,H_*) to biomass production flux (*J_B_*) at a fixed rubisco activity across a wide range of CA activities and Ci uptake rates. *J_L,tot_* was calculated as the sum of CO_2_ and HCO_3_^-^ leakage rates (*J_L,C_* + *J_L,H_*) with *J_L,C_* » *J_L,H_* in most conditions due to the much greater membrane permeability of CO_2_. So-called “futile cycling,” where leakage greatly exceeds biomass production (*J_L,tot_*/ *J_B_* » 1), occurs when CA and Ci uptake are co-expressed at extreme levels (redder colors). See *Supplementary Information* for detailed description of the co-limitation model.

When the HCO_3_^-^-dependence of biomass production was included in our model, it became possible to rationalize the growth improvements associated with heterologous expression of a CA or Ci transporter, as both of these activities can supply HCO_3_^-^ needed for biosynthesis (Figure 7A). When the modeled rubisco activity was low, growth was rubisco-limited: increasing rubisco activity (lighter colored lines in Figure 7B) or the CO_2_ concentration produced faster growth, but the growth rate was insensitive to CA activity because slow CO_2_ hydration (whether spontaneous or CA catalyzed) provided sufficient HCO_3_^-^ for the anabolic carboxylations involved in biomass production (Figures S11 and S12). When rubisco activity was set to higher values (darker lines in Figure 7B) the model entered a “bicarboxylation-limited” regime where increased rubisco activity did not affect the growth rate until CA activity was increased to supply HCO_3_^-^ (i.e. bicarbonate) for anabolic carboxylations. In bicarboxylation-limited conditions, additional HCO_3_^-^ could be supplied by CA, Ci uptake, or both (Figures S11). However, these activities are not equivalent, as we discuss below.

The co-limitation model also gave insight into the nature of the supposed “futile cycle,” which could now be framed quantitatively by comparing the leakage of inorganic carbon from the cell (flux *J_L,tot_* = *J_L,C_* + *J_L,H_*) to biomass production by rubisco and HCO_3_^-^-dependent carboxylases (*J_B_*, Fig. 7A). As CO_2_ is orders of magnitude more membrane-permeable than HCO_3_^-^ near neutral pH (16), CO_2_ leakage (*J_L,C_*) greatly exceeded the HCO_3_^-^ leakage flux (*J_L, H_*) in most conditions. Low Ci leakage is desirable; when *J_L,tot_* ≈ 0 the flux of Ci pumped into the cell is balanced by fixation of CO_2_ and HCO_3_^-^ such that no energy is “wasted” on Ci uptake. When CA (5) and Ci uptake (*χ*) activities were low, the model predicted slow growth and net diffusive Ci uptake (*J_L,C_, J_L,H_* < 0) to balance carboxylation (Figure 7C and S14). Increasing *χ* could cause *J_L,tot_* to change sign from negative (connoting passive uptake) to positive (connoting substantial leakage) passing through *J_L,tot_* = 0. If CA was also expressed - i.e. if *δ* was increased above a baseline value - *J_L,tot_* ≈ 0 could be achieved at lower *χ* values due to CA-catalyzed dehydration of imported HCO_3_^-^ producing CO_2_ used by rubisco. In other words, CA activity lowers the Ci uptake rate and, consequently, energy expenditure, required to achieve the same rate of biomass production (Figures 7C & S14). When *δ* and *χ* were both high, the model predicted substantial leakage, with *J_L,tot_* / *J_B_* ≈ 100 in extreme cases. Such extreme levels of futile cycling are likely incompatible with growth. If *χ* was low, in contrast, *δ* could be increased arbitrarily without incurring leakage because CAs do not pump Ci into the cell.

A second, less plausible way in which co-expression might improve growth is if CA and Ci transport are fast and membrane permeability to CO_2_ is far lower than typically assumed. In this case, the combination of Ci uptake and CA activity might form a CO_2_ pump that elevates C_in_ substantially above C_out_ to accelerate rubisco carboxylation (Figure S13). As HCO_3_^-^ is supplied sufficiently by Ci uptake in this case, the colimitation model predicted that increased rubisco fluxes would translate into faster growth. For this effect to arise, however, the membrane permeability to CO_2_ would have to be 100-1000 times lower than the measured (50) or calculated (51), so CO_2_ pumping is unlikely to explain the capacity of a CCMB1 to grow in ambient air when co-expressing Dab2 and Can (Fig. 6).

Taken together, our experiments and model helped outline a plausible trajectory for the co-evolution of the bacterial CCM with atmospheric CO_2_ levels. Presuming an ancestral autotroph with only a rubisco-driven CBB cycle and no CCM components, our data and model support a trajectory where CA and Ci transport are acquired together or serially (in either order) to support growth as atmospheric CO_2_ levels decreased (darker arrows in Fig. 1C). The order of acquisition might depend on the environmental pH, which strongly affects the extracellular HCO_3_^-^ concentration and, thereby, the expected efficacy of HCO_3_^-^ uptake (16). The co-limitation model helped us understand the potential advantages of expressing CA and Ci uptake together: modest co-expression can reduce energy expended on Ci pumping and balance the supply of CO_2_ and HCO_3_^-^ with the cellular demand for rubisco and bicarboxylation products (Figure 7C and S14). In CCMB1 we found that co-expression of CA and Ci transport supported growth in low CO_2_ environments (≈1%, Figure 4) and even permitted modest growth in atmosphere (Figure 6); cells expressing both activities would have been “primed” for the subsequent acquisition and refinement of proto-carboxysome structures that co-encapsulate rubisco and CA to enable robust growth at yet lower CO_2_ levels (e.g. ambient air, Figure 3). Notably, carboxysome shell proteins are structurally related to two ubiquitous protein families (65) and homologous to other metabolic microcompartments (28, 65), suggesting two plausible routes for the acquisition of carboxysome genes.

Results from our *E. coli* experiments indicated that such an evolutionary trajectory is “fitness positive” in that each step improves organismal fitness at some environmental CO_2_ level (Figs. 3, 4, and 6). Moreover, the fitness contribution of CA and Ci transport activities was only realized at intermediate pCO_2_ ≈ 1% in both *E. coli* (Fig. 4) and *C. necator* (Fig. 5), supporting the view that CO_2_ concentrations declining from high levels early in Earth history promoted the evolutionary emergence of bacterial CCMs.

### Concluding Remarks

There is great and longstanding interest in characterizing the composition of the atmosphere over Earth history (9, 11, 66). This interest is surely justified as the contents of the atmosphere affected the temperature, climate, and the chemical conditions in which life arose, evolved, and has been maintained. In the case of reactive species like O_2_, present-day measurements of old sedimentary rocks are quite informative: measurements of proxies for O_2_ reactivity in geological samples demonstrate that the Archean atmosphere contained very little dioxygen, with inferred O_2_ levels of 1 ppm or less (8, 9). Many posit that the Archean atmosphere also contained very high levels of CO_2_. However, since CO_2_ is far less reactive in Earth surface environments than O_2_, available geological constraints are meager and this inference stands on shakier ground.

One reason for presuming high CO_2_ levels in the early atmosphere is the paradox of the “Faint Young Sun,” which would not have been sufficiently luminous to permit liquid water on Earth without a very substantial greenhouse effect (67) or some other physical mechanism warming the Earth system (68). Other approaches to estimating ancient CO_2_ pressures rely indirectly on geochemical observations and arguments from the geological record. For example, some studies infer high Archean CO_2_ from the presence or absence of certain carbonate minerals in ancient surface environments (9), while others rely on models of biological CO_2_ fixation to estimate CO_2_ concentrations from the carbon isotope ratios of organic matter in sedimentary rocks (11, 12). These diverse approaches vary in the magnitude of their estimates, but each gives the impression that Archean CO_2_ greatly exceeded present-day levels of roughly 0.04% of a 1 bar atmosphere; indeed, some estimates of Archean CO_2_ pressures are as high ≈1 bar (9, 67). Silicate weathering coupled to the deposition of carbonate-bearing rocks represents a long-term crustal sink that — combined with subequal amounts of organic carbon burial — would have drawn a large reservoir of atmospheric CO_2_ down to pre-industrial levels (10, 69). Furthermore, modern plants, algae, and autotrophic bacterial lineages collectively evolved four distinct classes of CCM at least 100 times indicating that autotrophy has repeatedly and convergently adapted to decreasing environmental CO_2_ levels (13).

Carbon isotope methods for inferring historical changes in CO_2_ rely on uniformitarian models of biological carbon fixation to calculate atmospheric CO_2_ pressures from isotope ratio measurements of biogenic organic phases preserved in sediments and rocks (12). Yet autotrophy has undergone considerable evolution and diversification over the last ≈3 billion years, which complicates inference of ancient CO_2_ levels with models based on contemporary autotrophs. For example, rubiscos from different lineages display variable ^12^CO_2_ preferences (23, 70) and contemporary CCMs rely on Ci uptake systems that may fractionate C isotopes to a degree similar to rubisco (71). As demonstrated in the companion paper by Renee Wang and co-authors, the evolution of CCMs must be considered when attempting to reconstruct historical CO_2_ concentrations from the C isotope record.

Here we attempted to reconstruct the evolution of a bacterial CCM by constructing analogues of plausible ancestors. This investigation was initially motivated by a basic curiosity about the evolution of physiological complexity, hoping to resolve the apparent “irreducible complexity” of bacterial CCMs (Figure 1). Irreducible complexity is incompatible with evolution by natural selection, and so we hypothesized that ancestral CCMs conferred a growth advantage in the historical environments in which they arose, which we and others presume were characterized by high CO_2_ (5, 24–26).

Results from our experiments with rubisco-dependent *E. coli* and two bacterial autotrophs (*H. neapolitanus* and *C. necator)* supported this rationalization of CCM evolution by showing (i) that CO_2_ ≈ 1% improves the growth of “ancestral” CCMs (Figs. 3–6) and (ii) that all CCM genes are dispensable during growth in 5-10% CO_2_ (Fig. 2–5). This latter result suggested to us that atmospheric CO_2_ concentrations exceeded 1% of a 1 bar atmosphere when rubisco arose, which was likely more than 3 billion years ago (24, 72). If geological CO_2_ sinks later brought atmospheric CO_2_ to ≈1%, all organisms, including autotrophs, would have begun to evolve or acquire carbonic anhydrases and/or Ci transporters to provide the HCO_3_^-^ required for biosynthesis (Figure 7). An ancestral autotroph expressing both these activities may have had a growth advantage in relatively lower CO_2_ pressures < 1% (Fig. 6A), and would have been “primed” for the evolution of a CCM as the only missing component, the carboxysome, could have evolved from oligomeric host proteins (65) or adapted from a different metabolic microcompartment (28). Alternatively, it is possible that CAs (or Ci transporters) arose prior to rubisco and were already widespread at the time of rubisco evolution, in which case we might expect CO_2_ ≿ 1% when rubisco arose. Unfortunately, the convergent evolution of CA activity in several protein families (73) makes it very challenging to constrain the timing of CA evolution with comparative biological and molecular clock approaches; this issue concerns bacterial Ci transporters as well (17, 22).

It is intrinsically difficult to answer questions about Earth’s deep biological history; addressing such questions will surely require cooperation between scientific disciplines. Here, and in the companion paper by Renee Wang et. al., we took a “synthetic biological” approach to study the molecular evolution of bacterial CO_2_ fixation by constructing contemporary cells intended to resemble ancient ones. In both cases, our work highlighted the impact of environmental context (e.g. CO_2_ concentrations) and whole cell physiology (the requirement for HCO_3_^-^) on the evolution of CO_2_ fixation. That is, neither rubisco nor the CCM should be considered in isolation, but rather in the context of a metabolism that demands both CO_2_ and HCO_3_^-^ to produce biomass (55–61, 71). We hope that future research advances the synthetic approach to studying evolution and fully expect that this approach will enrich our understanding of biological processes that have shaped the evolution of biogeochemical cycles on Earth.

## Methods

### Strains, plasmids and genomic modifications

Strains and plasmids used in this study are documented in Supplementary Tables S4 and S5. The rubisco-dependent *E. coli* strain CCMB1 was derived from *E. coli* BW25113 and has the genotype BW25113 *ΔrpiAB Δedd ΔcynT Δcan,* as documented in (4). To construct CA deficient mutants of *C. necator* H16, we first knocked out the hdsR homolog *A0006* as removal of this restriction enzyme increases electroporation efficiency (41). The double CA knockout, *C. necator* H16 Δ*A0006 Δcan Δcaa,* was constructed by repeated rounds of selection and counter-selection by integrating a construct encoding both kanamycin resistance and *sacB* for counter-selection. Protocols for plasmid construction and genomic modification of *C. necator* are fully described in the *Supplementary Information.*

### Genome-wide fitness measurements in *H. neapolitanus*

Competitive fitness assays were performed following (22). The barcoded *H. neapolitanus* transposon library generated for that work was thawed and used to inoculate three 33 ml cultures that were grown overnight at 30 C in DSMZ-68 with 10 μg/ml Kanamycin. These pre-cultures were grown to an OD ≈ 0.07 (600 nm) in 5% CO_2_. The library was subsequently back-diluted 1:64 and grown in various CO_2_ concentrations (10%, 5%, 1.5%, 0.5% and ambient CO_2_) on a platform shaker (New Brunswick Scientific Innova 2000, 250 RPM) in a Percival Intellus Incubator; 20 ml of pre-culture was pelleted by centrifugation (15 min at 4000g) and saved as a T0 reference. Upon reaching 6.5-7.5 doublings, 50 mls of culture was spun down and gDNA was extracted for barcode PCRs as described in (6); barcodes were previously mapped to the genome via TnSeq (22). PCRs were purified (Zymo Research Clean and Concentrator kit) and pooled for sequencing on an Illumina MiSeq with 150 bp single end reads. We used the software pipeline from (32) to analyze barcode abundance data. Briefly, fitness of individual mutant strains was calculated as the log_2_ of the ratio of barcode abundance in the experimental condition over abundance in the T0. Gene-level fitness values were then calculated considering all transposon insertions expected to disrupt an individual gene. Each CO_2_ concentration was tested in biological duplicate except for 5% CO_2_, which was assayed in biological quadruplicate.

### *E. coli* growth conditions

*E. coli* strains were grown at 37 °C. 60 μg/ml kanamycin and 25 μg/ml chloramphenicol were used for selection during routine cloning and propagation. For strains carrying two selectable markers, antibiotics were used at half concentration (30 μg/ml kanamycin, 12.5 μg/ml chloramphenicol). All CCMB1 cultures were grown in 10% CO_2_ unless otherwise specified. When rubisco-independent growth was desired, CCMB1 was propagated in rich LB media. CCMB1 was cultured in a rubisco-dependent manner in M9 minimal media supplemented with trace elements and 0.4% glycerol (v/v) as described in (4). Experiments presented in Figures 3, 4 and 6 were conducted in 96-well plates in a gas-controlled plate reader (Tecan Spark). For the data in Figure 3, 100 nM anhydrous tetracycline (aTc) was supplied to induce expression from p1A, pCB’ and pCCM’ plasmids. For the data presented in Figures 4 and 6, a constitutive version of p1A was used (p1Ac) and aTc was omitted from pre-cultures. These strains also carried a dual-expression plasmid, pFC, for inducible expression of the DAB2 Ci transport operon and the *can* carbonic anhydrase. 100nM aTc was supplied to induce DAB2 and 1 mM IPTG for *can*.

Pre-cultures were inoculated into 5 ml of M9 glycerol with 30 μg/ml kanamycin, 12.5 μg/ml chloramphenicol and appropriate induction. 1 ml of each culture was transferred to a separate tube, which was incubated in ambient CO_2_ as a negative control; the remaining 4 ml was incubated in 10% CO_2_. CCMB1 strains carrying active site mutants of rubisco (cbbL K194M) were pre-cultured in LB with the same antibiotic concentrations. Once cultures reached saturation they were centrifuged at 4000 g for 8 min. Pellets were washed in 10 ml of M9 with no carbon source (M9 NoC) and resuspended in 5 ml M9 NoC. Optical density was measured in five-fold dilution at 600 nm (Genesys 20 spectrophotometer, Thermo Scientific) and cultures were normalized to 0.5 OD units. 96-well plates were inoculated by adding 2.5 μl of OD-normalized preculture to 247.5 μl M9 glycerol supplemented with appropriate antibiotics and induction. Kanamycin was omitted as the plasmid carrying kanamycin resistance also expresses rubisco, which is required for growth in glycerol media (4). To minimize evaporation during multi-day cultivations, 150 μl sterile water was added to the reservoirs between the wells and the plate was incubated inside of a small humidity cassette (Tecan) with 3 ml sterile water added to each reservoir. Plates were incubated with shaking for at least 4 days in a Tecan Spark plate reader configured to control the CO_2_ concentration. The humidity cassette was replenished after 48 hours. All experiments were performed in biological quadruplicate and technical triplicate.

### *C. necator* growth conditions

*C. necator* strains were grown in ambient CO_2_ unless otherwise specified, and 200 μg/ml kanamycin was added to select for plasmid retention. When rubisco-independent heterotrophic growth was desired, *c. necator* was cultured in LB at 30 °C. For autotrophic growth experiments, strains were pre-cultured in 5 mL of LB media in 10% CO_2_ in 25 mL tubes with 20 mm butyl stoppers sealed by aluminum crimping. Precultures were incubated for two days at 30 °C with 200 rpm shaking, washed three times in *Cupriavidus* minimal media, and inoculated to an OD of 0.1 (550 nm) in minimal media in a 165 mL flask with a 20 mm butyl stopper sealed by aluminum crimping. *Cupriavidus* minimal growth media contained 3.24 mM MgSO_4_, 0.42 mM CaCl_2_, 33.5 mM NaH_2_PO_4_, 32.25 mM Na_2_HPO_4_, 2.6 mM K_2_SO_4_, 1 mM NaOH, 1.87 mM NH_4_Cl, and 1 mL/L of a 1000x trace minerals solution containing 480 mg/L CuSO_4_·5H_2_O, 2.4 g/L ZnSO_4_·7H_2_O, 2.4 g/L MnSO_4_·H_2_O, and 15 g/L FeSO_4_·7H_2_O (pH 6.7). The flasks were evacuated and the headspace was filled with 60% H_2_, 10% O_2_, and indicated concentrations of CO_2_. The balance was atmospheric air. Cells were grown for 48 hours at 30° C with 200 rpm shaking, and OD550 values were taken using a Molecular Devices SpectraMax M2 spectrophotometer.

## Acknowledgements

We dedicate this paper to the memory of Danny Salah Tawfik (z”l), a luminary to the scientific community and a dear friend and mentor to AIF. Danny’s many studies of enzyme evolution taught us that evolutionary timescales are accessible in carefully-designed lab-scale experiments and his active cultivation of relationships that transcend generational, professional and cultural divides continues to inspire. We are also indebted to Arren Bar-Even (z”l) for formative conversations guiding this work. Many thanks to Darcy McRose, Elad Noor, and Renee Wang for extensive comments on the manuscript and to Cecilia Blikstad, Julia Borden, Vahe Galstyan, Josh Goldford, Ron Milo, Robert Nichols, Naiya Phillips, Noam Prywes, and Gabe Salmon for helpful input. This research was supported in part by an NSF Graduate Research Fellowship (to A.I.F), NSF Grant No. PHY-1748958, the Gordon and Betty Moore Foundation Grant No. 2919.02, and the Kavli Foundation (to A.I.F), the US Department of Energy (DE-SC00016240 to D.F.S.) and Royal Dutch Shell (Energy and Biosciences Institute project CW163755 to D.F.S. and S.S.).

