## Supplementary Text and Figures for "Trajectories for the evolution of bacterial CO_2_-concentrating mechanisms"

**This PDF file includes:**

Supplementary text
Figures S1 to S14
SI References

**Other supplementary materials for this manuscript include the following:**

Supplementary Tables S1-S5

**Supplementary Methods** 3
Plasmid Construction 3
Electroporation of CCMB1 E. coli 3
Manipulation of the C. necator genome 3
Plasmid transformation of C. necator 4
Modeling the co-limitation of autotrophic growth 5
Carbonic anhydrase cannot reasonable act as a CO2 pump alone 5
A model of autotrophy including the HCO3--dependence of growth 7
Choosing realistic ranges for parameter values 8
On the requirement for bicarbonate for biosynthesis 9
**Supplementary Figures** 12
**Supplementary References** 26

### Supplementary Methods

#### Plasmid Construction

Genes of interest were amplified by PCR and cloned into their respective vectors using Gibson Assembly. Plasmids were then transformed into chemically competent NEB Turbo *E. coli* cells in most cases. Single colonies were inoculated into 5-8 mL LB media with appropriate antibiotics and mini-prepped once turbid (Qiagen QIAPrep spin kit). For the construction of p1Ac, which constitutively expresses *prk*, we used CCMB1 as the cloning strain (1). In addition, to ensure that *prk* expression was not deleterious, as it is for wild-type *E. coli* (2), CCMB1:p1Ac transformants were cultured in a *prk*-dependent manner M9 glycerol media. Plasmid sequences were verified by Sanger sequencing at the UC Berkeley DNA sequencing facility. Details of plasmids used in this study are documented in Table S5 and plasmids have been deposited to Addgene at https://www.addgene.org/David_Savage/.

#### Electroporation of CCMB1 *E. coli*

Electrocompetent CCMB1 stocks were prepared by standard methods from cultures grown in LB media under 10% $CO_2$. Plasmids were transformed via electroporation with the following protocol. A 50 µl aliquot of electrocompetent CCMB1 was placed on ice until thawed. 100 ng of mini-prepped plasmid (100 ng each if a double transformation) was then added, gently mixed, and left to incubate for 10 minutes. The transformation aliquot was subsequently transferred to a chilled 1mm cuvette (Biorad Gene Pulser) and pulsed in a Gene Pulser Xcell Microbial System electroporator (1800 V, 200Ω, 25µF). 500 µl SOC was added to the cuvette and the resulting culture was pipetted into a 14 ml round-bottom falcon tube and placed in 10% $CO_2$ to incubate for 1 hour. Do not recover in ambient $CO_2$ or in a microcentrifuge tube, as it is important to recover CCMB1 strains in high $CO_2$ to ensure that recovery is independent of the transformed plasmid(s). After incubation, 200 µl of the culture was plated on an LB agar plate with appropriate antibiotics. When preparing S17 *E. coli* donor cells for conjugation with *C. necator*, plasmids were transformed by the same method, except the recovery was done in ambient $CO_2$ for 30 minutes.

#### Manipulation of the *C. necator* genome

The triple knockout mutant *C. necator* Δ*A0006*Δ*can*Δ*caa* was produced by iterative rounds homologous recombination (to generate a desired mutation) followed by *sacB* counterselection to cure the kanamycin resistance marker integrated at the target locus (3). Homologous recombination was achieved by conjugation with *E. coli* S17 carrying a mobilizable vector encoding 500 bp homology arms flanking a cassette encoding kanamycin resistance and *sacB* counter selection. For each individual knockout, a pKD19-mobSacB plasmid was generated with 500 bp homology arms directly flanking the target gene. This plasmid was transformed into *C. necator* by conjugation with *E. coli* S17 and plated onto LB agar supplemented with 200 µg/ml kanamycin to select for integrants and 10 µg/ml gentamicin to select against residual *E. coli*.

Single integrant colonies were inoculated into LB with 10 µg/ml gentamicin and 20 µg/ml kanamycin and incubated in 30 °C until turbid. Genomic integration was verified by colony PCR using a primer set where one primer annealed to the genome and the other primer annealed to the plasmid backbone. Verified colonies were inoculated into salt-free LB (10 g/L tryptone, 5 g/L yeast extract) supplemented with 10 µg/ml gentamicin and 100 mg/ml sucrose and incubated at 30 °C for 48-72 hours to select against *sacB* activity. Strains were then streaked on two different LB plates: one without NaCl, but containing 10 µg/ml gentamicin and 50 mg/ml sucrose and a second plate with NaCl, 10 µg/ml gentamicin and 200 µg/ml kanamycin. Colonies that grew on sucrose but not on kanamycin were genotyped by colony PCR using a pair of primers that annealed upstream and downstream of the target gene. PCRs were run on an agarose gel to ensure prospective knockouts were not wild-type revertants. The final strain, *C. necator* Δ*A0006*Δ*can*Δ*caa* was further verified by phenotype: it fails to grow heterotrophically in ambient air, but is able to grow under elevated $CO_2$ (4, 5).

### Plasmid transformation of *C. necator*

To enable routine electroporation of plasmids into *C. necator* H16, we first knocked out the *hdsR* homolog *A0006* as removal of this restriction enzyme increases electroporation efficiency (3, 6). Electrocompetent stocks of *C. necator* Δ*A0006*-derived strains (including the various knockouts) were made according to a protocol from (3) with the following modifications. A colony of the strain was inoculated into LB with 10 µg/ml gentamicin. Once turbid, the pre-culture was added to 100 mL fresh media and let grow until it reached an OD600 between 0.6-0.8. ΔA0006 was grown in ambient $CO_2$ and ΔA0006Δ*can*Δ*can* was grown in 10% $CO_2$. Cells were then chilled, shaking in an ice slurry until they reached 4 °C. The culture was split into two 50 ml Falcon tubes and centrifuged at 4000*g* for 10 minutes at 4 °C. The supernatant was decanted and pellets were washed twice with 50 ml ice cold sterile water and once with 50 ml 10% glycerol. The pellets were then resuspended in 0.75 ml 10% glycerol, pooled, and 100 ul aliquots were flash-frozen in liquid nitrogen for storage at -80 °C.

For plasmid transformation, a 100 µl aliquot of *C. necator* was thawed on ice. Upon thawing, 500 ng of plasmid was added, gently mixed, and left to incubate on ice for 5 minutes. The aliquot was then transferred to a 1 mm electroporation cuvette (Biorad Gene Pulser) and pulsed in a Gene Pulser Xcell Microbial System electroporator (2300 V, 200Ω, 25µF). The sample was then immediately resuspended in 1 ml of LB supplemented with 10 mg/ml fructose, transferred into a 14 ml round-bottom falcon tube, and recovered in a 30 °C for 2 hours (H16 Δ*A0006* in ambient $CO_2$, H16 Δ*A0006*Δ*can*Δ*caa* in 10% $CO_2$). 200 µl was then plated on LB agar plates with 10 µg/ml gentamicin, 200 µg/ml kanamycin, and 10 mg/ml fructose and placed in a 30 °C incubator at ambient $CO_2$ or 10% $CO_2$ (depending on the strain) for 48 hours.

### Modeling, data and analysis

The dual-limitation model was elaborated in Mathematica 12 (Wolfram) and steady-state solutions were translated to Python for further analysis and plotting. All data analysis was performed using Python 3.8 and Jupyter notebooks. Data and code required to generate all figures is available at https://github.com/flamholz/ccm_evolution.

### Modeling the co-limitation of autotrophic growth

#### Carbonic anhydrase cannot reasonably act as a $CO_2$ pump alone

Our model considers an autotroph with no CCM that uses rubisco to fix $CO_2$ in an environment with fixed extracellular $CO_2$ and $HCO_3^-$ concentrations, $C_{out}$ and $H_{out}$. We further assume that these extracellular species are in equilibrium with respect to the pH, i.e. that $H_{out}/C_{out} = K_{EQ}(pH)$, and that the intracellular pH is the same as the extracellular pH so that the pH-dependent equilibrium constant $K_{EQ}(pH)$ is equal on both sides of the cell membrane. This assumption of equal pH equilibrium is not required but simplifies the model (7). We now write differential equations describing the time evolution of the intracellular $CO_2$ and $HCO_3^-$ concentrations, $C_{in}$ and $H_{in}$, at first ignoring the $HCO_3^-$ dependence of growth to illustrate that it must be included.

Since $CO_2$ and $HCO_3^-$ have diffusion constants of ≈ $10^3$ $\mu m^2/s$ corresponding to diffusion timescales of $R^2/6D \approx 10^{-4}$ s over the ≈ 1 micron lengths of bacterial cells, we will assume that their concentrations are spatially homogeneous inside and outside the cell (8). We also assume all enzyme-catalyzed reactions have first-order kinetics, i.e. substrate concentrations are substantially lower than Michaelis constants ($[S] \ll K_M$). These assumptions give the following equations:

$$\frac{dC_{in}}{dt} = \alpha(C_{out} - C_{in}) - \gamma C_{in} - (\delta C_{in} - \phi H_{in})$$

$$\frac{dH_{in}}{dt} = \beta(H_{out} - H_{in}) + (\delta C_{in} - \phi H_{in})$$

Here we treat both $CO_2$ and $HCO_3^-$ as entering the cell passively with “effective permeabilities” α and β. These effective permeabilities account for the surface area to volume ratio of bacterial cells, which, for rod shaped cells around the size of *E. coli*, is $SA/V \approx 4\ \mu m^{-1}$ (BNIDs 101792 and 114924) as we discuss below.

$\gamma C_{in}$is a linearized expression for rate of irreversible $CO_2$ fixation by rubisco, where $\gamma = k_{cat}[rubisco]/K_M$ assuming a Michaelis-Menten formalism and $C_{in} \ll K_M$. In contrast to rubisco, the CA reaction is reversible. As such, $(\delta C_{in} - \phi H_{in})$ the balance of the rates of $CO_2$ hydration $(\delta C_{in})$ and $HCO_3^-$ dehydration $(\phi H_{in})$, assuming each of these reactions are in their linear regimes as well. While the assumption of linearity is not required, it is also not counterfactual: typical $K_M$ values measured for bacterial rubiscos (9) and carbonic anhydrases (10) are comparable to equilibrium concentrations of $CO_2$ and $HCO_3^-$ in water in equilibrium with ambient air at 25 C (Figure S9).

We set both derivatives to 0 and solve for the steady-state values of $C_{in}$ and $H_{in}$.

$$C_{in} = \frac{C_{out}(K_{EQ}\beta\phi + \alpha(\beta + \phi))}{\beta(\alpha + \gamma + \delta) + \phi(\alpha + \gamma)}$$

$$H_{in} = \frac{C_{out}(\alpha\delta + K_{EQ}\beta(\alpha + \gamma + \delta))}{\beta(\alpha + \gamma + \delta) + \phi(\alpha + \gamma)}$$

If we further assume that CA activity is negligible, i.e. that $\delta, \phi \approx 0$, then we recover the solution from our simplified main-text calculation where $C_{in} = \frac{C_{out}\alpha}{\alpha+\gamma}$ is independent of $H_{in}$. As a reminder, we used this equation to calculate that $C_{in} > 0.9C_{out}$ in the absence of CA activity, even when rubisco comprises 20% of total protein.

The above calculation implies that CA expression could increase $C_{in}$ by at most 10% because CAs are not coupled to any energy source and, therefore, cannot increase $C_{in}$ above $C_{out}$. This calculation depends, of course, on the rubisco kinetics and expression ($\gamma$) and membrane permeability to $CO_2$ ($\alpha$). Rubisco kinetics have been studied in great depth and are well-constrained (9, 11). Similarly, many generations of physical chemists have studied the permeability of lipid membranes to small molecules and developed theory to estimate membrane permeabilities (12–14). Nonetheless, membrane permeabilities can depend on the lipid composition of the membrane and the complement of protein channels embedded therein (15).

Assuming that rubisco fixation is the sole growth-limiting reaction, we can estimate the exponential growth rate from $C_{in}$ by calculating the rubisco fixation rate $\gamma\ C_{in} \approx 9\text{x}10^3$ µM/s. Here we took $C_{out} \approx 10$ µM, which is roughly Henry's law equilibrium with present-day atmosphere at 25 °C, $\alpha = 10^4\ s^{-1}$ and $\gamma = 10^3\ s^{-1}$. We expound on this choice of values in the main text and below. Assuming a cell volume of ≈ 1 fL (BNIDs 104843, 100004), $9\text{x}10^3$ µM/s equals a fixation rate of roughly $5\text{x}10^6\ CO_2$/s or ≈ $10^{10}\ CO_2$/hr. An *E. coli* cell of this volume contains ≈$10^{10}$ carbon atoms (BNID 103010) and Cyanobacteria do not differ substantially from *E. coli* in carbon content (compare BNIDs 105530 and 111459). Therefore, assuming no loss of fixed carbon, such a Cyanobacterium would double once an hour. Autotrophic respiration, which equals the difference between gross and net fixation, is typically less than 50% of gross both in pure cyanobacterial cultures (16) and natural ecosystems (17) implying a doubling time of at most 2 hours.

Given the model articulated above, a 10% increase in $C_{in}$ (e.g. due to CA expression) can increase the rubisco carboxylation rate by at most 10%. As rubisco is required for producing all biomass carbon in autotrophy, a 10% increase in the rate of rubisco carboxylation can increase the exponential growth rate by at most 10%. However, in Figures S5-6 the "rubisco alone" strain did not meaningfully grow in 0.5% $CO_2$ while the strains expressing a CA or Ci transporter grew robustly. These qualitative effects indicated that we should look for a mechanism that can improve growth by more than ≈10%. As described in the main text and the following section, the cellular demand for $HCO_3^-$, which is required for several anabolic carboxylation reactions (18–21), is one such mechanism.

Notably, $CO_2$ and $HCO_3^-$ do interconvert spontaneously. The spontaneous reaction is associated with relatively slow kinetics, with $\delta_{spont} \approx 10^{-2}\ s^{-1}$ and $\phi_{spont} \approx 4 \times 10^{-3}\ s^{-1}$ near pH 7 (7, 22). Therefore, zero CA expression does not entail our above assumption that $\delta, \phi \approx 0$. Rather, to recover the expression for $C_{in}$ above, we require that $\phi_{spont} \ll \beta$, $K_{EQ}\phi_{spont} \ll \alpha$, and $\delta_{spont} \ll \gamma = k_{cat}[rubisco]/K_M$. The latter is true for any modest level of rubisco expression: as typical rubiscos have $k_{cat}/K_M \approx 10^5\ M^{-1}s^{-1}$ (9) and a bacterial rubisco should have a concentration of at least $10^{-6}$ M (23), $\gamma \geq 10\ s^{-1} \gg \delta_{spont}$ (Fig. S9). Similarly, $\beta = SA \times P_H/V \approx 10^{-2}\ s^{-1}$ is roughly five times larger than $\phi_{spont}$ near pH 7 (7). Finally, near pH 7, $K_{EQ} \approx 10$ and so $K_{EQ}\phi_{spont} \approx 4 \times 10^{-2}\ s^{-1}$. This value is similar in scale to $\beta$, which is 2-3 orders smaller than $\alpha$ (7). Therefore, the simplified equation above is supported near pH 7.

#### A model of autotrophy including the $HCO_3^-$-dependence of growth

Our above calculation indicated to us that CA cannot act as a $CO_2$ pump and that, therefore, some factor is missing from the naive model of autotrophy given above. We assume that the missing factor is the ubiquitous dependence of microbial growth on $HCO_3^-$. This dependence is well-documented for heterotrophic microbes and stems from the use of bicarbonate-dependent carboxylases in nucleotide, amino acid, and lipid biosynthesis (18–20). Similar dependencies have been observed in land plants (21) and manually-curated metabolic models of autotrophs include these same reactions (21, 24, 25), indicating that this dependence of growth on $HCO_3^-$ is very widespread, perhaps even universal. Furthermore, our data demonstrate that the model bacterial chemolithoautotroph, *C. necator*, depends on CA for robust growth in relatively low $CO_2$ levels (1.5% or lower, Fig. 5B). As this growth defect is complemented by expression of the DAB2 Ci transporter (Fig 5B), we interpret these data as supporting the hypothesis that *C. necator* depends on $HCO_3^-$ for autotrophic growth in ambient air.

We therefore augmented our model to reflect the apparent ubiquity of bicarbonate dependence by including (i) an $HCO_3^-$ consuming flux, $-\omega H_{in}$, representing bicarbonate-dependent carboxylation in central metabolism (henceforth "bicarboxylation") and (ii) a flux, $+\chi H_{out}$, producing intracellular $HCO_3^-$ representing energized bicarbonate uptake systems like the DABs (26) or Cyanobacterial *sbtA* transporters (27, 28).

$$\frac{dC_{in}}{dt} = \alpha(C_{out} - C_{in}) - \gamma C_{in} - (\delta C_{in} - \phi H_{in})$$

$$\frac{dH_{in}}{dt} = \beta(H_{out} - H_{in}) + (\delta C_{in} - \phi H_{in}) + \chi H_{out} - \omega H_{in}$$

Following the example of the Farquahar model of photosynthesis (29), we assume that the flux to biomass $J_B$ is determined as the minimum of two fluxes: the $CO_2$-dependent flux through rubisco ($\gamma C_{in}$) and flux through $HCO_3^-$ dependent carboxylation reactions ($\omega H_{in}$). This is co-limitation expressed as $J_B = \min(\gamma C_{in}, \omega H_{in} / q)$ where $q$ is the fraction of biomass carbon deriving from $HCO_3^-$. The exponential growth rate $\lambda$ can be estimated from $J_B$ by noting that a typical bacterial cell contains $\approx 10^{10}$ carbon atoms (BNID 103010). For simplicity we ignore the carbon cost of cellular maintenance, though this could be included in future renditions of the model.

Steady-state solutions are given below. These values determine the steady-state rates of rubisco carboxylation and bicarboxylation, which, in turn, determines the biomass production flux and exponential growth rate.

$$C_{in} = \frac{C_{out}(K_{EQ}\phi(\beta + \chi) + \alpha(\beta + \phi + \omega))}{\beta(\gamma + \delta) + \gamma\phi + \omega(\gamma + \delta) + \alpha(\beta + \phi + \omega)}$$

$$H_{in} = \frac{C_{out}(\alpha\delta + K_{EQ}(\alpha + \gamma + \delta)(\beta + \chi))}{\beta(\gamma + \delta) + \gamma\phi + \omega(\gamma + \delta) + \alpha(\beta + \phi + \omega)}$$

It is evident from these expressions that the rate of biomass production $J_B = \min(\gamma C_{in}, \omega H_{in} / q)$ will depend on $H_{in}$ in some circumstances and on $C_{in}$ in others. For example, if we assume $\delta, \phi, \chi \approx 0$, we find $H_{in} = C_{out} K_{eq} \beta / (\beta + \omega)$. Therefore, if CA and Ci uptake activities are negligible and the $HCO_3^-$ permeability $\beta$ is much smaller than the bicarboxylaton activity $\omega$, $H_{in}$ will be small and growth will be limited by low bicarboxylation flux. In the following sections we describe how we set reasonable ranges for all model parameters in order to examine the dependence of autotrophic biomass production on the activity of rubisco, CA, and Ci uptake systems.

### Choosing realistic ranges for parameter values

We assume the pH is the same both inside and outside the cell for simplicity. Furthermore, we choose pH 7.1 since the effective $pK_a$ between $CO_2$ and $HCO_3^-$ is roughly 6.1 in biological salt concentrations (see supplement of (7) for detail). According to the Henderson-Hasselbalch relation, $pH = pK_a + \log_{10}([HCO_3^-]/[CO_2])$, so the choice of pH = 7.1 sets the equilibrium constant $K_{EQ}(pH) = [HCO_3^-]/[CO_2] = 10^1$ both inside and outside the cell (7).

Since we endeavor to explain phenotypes observed in relatively low $CO_2$ levels (e.g. ambient air in Fig. 6 and 0.5-1.5% $CO_2$ in Figs. 4-5), we assume the extracellular $CO_2$ concentration is in Henry's law equilibrium with present day atmosphere (≈0.04% $CO_2$). This gives $C_{out} \approx 15$ µM (7, 30) and, with $K_{EQ} = 10$, $H_{out} = 150$ µM. For the permeability of the cell membrane to $CO_2$ and $HCO_3^-$, we use $P_C = 3\times10^3$ µm/s and $P_H = 10^{3.2-pH} \times 30$ µm/s ≈ $4\times10^{-3}$ µm/s following (7). The latter relation calculates the permeability of $HCO_3^-$ from its pH-dependent abundance and the permeability of $H_2CO_3$, assuming that $HCO_3^-$ has negligible permeability when compared to $H_2CO_3$ due to its charge. This calculation is described in detail in the supplement of (7). We multiply these permeabilities by the surface area to volume ratio SA/V ≈ 4 $\mu m^{-1}$ to obtain estimates of $\alpha \approx 1.2 \times 10^4\ s^{-1}$ and $\beta \approx 1.6 \times 10^{-2}\ s^{-1}$.

We are left to choose ranges for the enzymatic activity parameters $\gamma, \delta, \phi, \omega$ and $\chi$. First, we note that the the CA activity parameters $\delta$ and $\phi$ must be must be consistent with the equilibrium constant $K_{EQ}$ (i.e. must obey the Haldane relation). If the CA reaction was allowed to equilibrate, it would carry no net flux and $\delta C_{in} - \phi H_{in} = 0$. In these conditions $K_{EQ} = \frac{H_{in}}{C_{in}} = \frac{\delta}{\phi}$, giving $\phi = \frac{\delta}{K_{EQ}}$.

To set ranges for enzyme activities $\gamma$ (rubisco carboxylation) and $\delta$ ($CO_2$ hydration by CA), we reviewed literature values for $k_{cat}/K_M$ for rubiscos (9) and CA (10, 31). The geometric mean of measured rubisco $k_{cat}/K_M$ values is ≈ 0.2 $\mu M^{-1}\ s^{-1}$ with a multiplicative standard deviation of roughly two-fold (Figure S9). A typical protein concentration might range between 0.1 and 100 µM (23). As rubisco is typically one of the most abundantly expressed proteins in autotrophic cells (32), we extend this range to 0.1 µM - 1 mM implying that $\gamma$ ranges from $\approx 10^{-2}$-$10^3\ s^{-1}$. Note that we are using µM units for both the enzyme and substrate so that $\gamma C_{in}$ has units of µM/s carbon consumed. For CA, the geometric mean $k_{cat}/K_M$ value in the direction of $CO_2$ hydration is ≈ 20 $\mu M^{-1}\ s^{-1}$ with a multiplicative standard deviation of roughly seven-fold (Figure S9). CA is not typically as highly-expressed as rubisco, so a plausible range for $\delta$ is perhaps 0.1-

$10^4$ $s^{-1}$ when a CA is expressed. As noted above, the spontaneous reaction is characterized by $\delta_{spont} \approx 10^{-2}\, s^{-1}$.

When environmental $CO_2$ concentrations are sufficiently high, rubisco and CA can become $CO_2$ saturated and our assumption of linear kinetics is violated as enzymatic rates become zero order (i.e. independent of substrate concentrations $C_{in}$ and $H_{in}$). This can be addressed by a simple modification of the model, setting the rubisco rate to $k_{cat}$ [*rubisco*] and the CA rate to ($k_{cat,H}$ - $k_{cat,D}$)[*CA*] as appropriate. Here $k_{cat,D}$ is the $k_{cat}$ in the direction of $CO_2$ hydration and $k_{cat,D}$ is calculated from $k_{cat,H}$ via the Haldane relation as described above. This latter relation supposes that CA is substrate-saturated in both hydration and dehydration directions, i.e. saturated by $CO_2$ and $HCO_3^-$ both. When such a model is appropriate, realistic $k_{cat}$ values are required. Figure S10 shows that rubisco $k_{cat}$ values range from roughly 1-10 $s^{-1}$ (geometric mean 3.3 $s^{-1}$ with a multiplicative standard deviation of 1.5 fold) and $k_{cat}$ values for CA-catalyzed $CO_2$ hydration range from ≈$10^4$-$10^6$ $s^{-1}$ (geometric mean $1.3 \times 10^5$ $s^{-1}$ with a multiplicative standard deviation of 6.4 fold).

Only $\omega$ and $\chi$ remain to be set. We chose $\omega = \gamma / q$ with $q$ = 100 to reflect our assumption that both rubisco and bicarboxylation processes contribute to biomass production in a roughly fixed proportion ($q$), but that rubisco is responsible for the production of nearly all biomass carbon in autotrophy (we assume 99%) and bicarboxylation is responsible for the remainder (1%). We used the same value of $q$ in calculating the biomass flux from the principle of co-limitation, i.e. $J_B = \min(\gamma\, C_{in}, \omega H_{in} / q)$. This amounts to assuming that the cell regulates the bicarboxylation and rubisco capacities to match their relative contributions to biomass production. Our assumption that $\omega$ is proportional to $\gamma$ can be omitted, but this yields a model with an additional free parameter that is challenging to constrain from data.

To set $\chi$, we consider measurements of saturated Ci uptake rates in Cyanobacteria, which are on the order of 10-100 µmol per mg chlorophyll per hour (33). Since a typical cyanobacterial cell contains ≈$10^{-11}$ mg chlorophyll (34), the per-cell rates are at most $10^{-9}$ µmol/hour, or $3 \times 10^{-13}$ µmol/s into a volume of ≈1.5 µm$^3$ = $1.5 \times 10^{-15}$ L. Uptake rates in this range would contribute ≈ +200 µM/s to $dH_{in}/dt$. If $\chi\, H_{out} \leq 200\ \mu M/s$ and $H_{out} = 100 - 2000\ \mu M$ depending on the pH then $\chi \leq 2\ s^{-1}$. Note that Figure 7 and S11-14 use wider ranges for $\gamma$, $\delta$ and $\chi$ than calculated here in order to illustrate the behavior of the model with two-dimensional plots.

### On the requirement for bicarbonate for biosynthesis

One way to examine the role of bicarbonate dependent carboxylation in our model is to set the bicarboxylation rate constant $\omega$ = 0. This gives

$$C_{in} = \frac{C_{out}(\alpha(\beta + \phi) + K_{EQ}\phi(\beta + \chi))}{\beta(\alpha + \gamma + \delta) + \phi(\alpha + \gamma)}$$

$$H_{in} = \frac{C_{out}(\alpha\delta + K_{EQ}(\alpha + \gamma + \delta)(\beta + \chi))}{\beta(\alpha + \gamma + \delta) + \phi(\alpha + \gamma)}$$

We see that $C_{in}$ and $H_{in}$ remain interdependent, i.e. the processes that produce $H_{in}$ like $CO_2$ hydration by carbonic anhydrase ($\delta$) and active Ci uptake ($\chi$) are represented in the equation for $C_{in}$ and vice versa. Nonetheless, these processes have negligible effect on $CO_2$ fixation by rubisco because (i) $C_{in}$ uniquely determines the rubisco rate in our model, and (ii) literature values for $CO_2$ permeability are high enough that (iii) rubisco cannot reduce $C_{in}$ much beneath $C_{out}$, as described above. Figures S12 and S14 illustrate this point by showing that order-of-magnitude changes to $\delta$, $\chi$ and $\gamma$ do not substantially affect $C_{in}$. In particular, in Figure S12A, $C_{in} \approx C_{out}$ until rubisco activity reaches very high levels $\gamma \approx 10^4$ $s^{-1}$. This is a simple consequence of the fact that the measured $CO_2$ permeability of biological membranes ($\alpha$) is quite high. Figure S13 illustrates this point using a model with substantial Ci uptake activity ($\chi$ = 100 $s^{-1}$) and an unrealistically low value of $\alpha$ = 12 $s^{-1}$ (1000-fold smaller than we estimated above). Very low $\alpha$ values enable CA and Ci uptake to act in concert to pump $CO_2$ into the cell by (i) actively taking up $HCO_3^-$, and (ii) converting $HCO_3$ into $CO_2$ via CA, which is (iii) retained in the cell when the membrane permeability to $CO_2$ ($\alpha$) much smaller than calculated or measured (12, 14).

If order-of-magnitude changes to $\delta$ and $\chi$ do not affect $C_{in}$ (when realistic $\alpha$ values are used), then the rubisco carboxylation flux cannot change and we must invoke another mechanism to explain the observed phenotypes. As discussed in the main-text and above, we assumed that the ubiquitous requirement for $HCO_3^-$ as the substrate for biosynthetic carboxylases is the underlying mechanism. Once we described the growth rate as mathematically coupled to both rubisco carboxylation of $CO_2$ and biosynthetic carboxylation of $HCO_3^-$, we found that changes in CA and Ci uptake activities do produce changes in growth (Figure S12).

### A quantitative view of futile cycling

Figures 7C and S14 document the effects of simultaneously varying CA activity ($\delta$) and Ci uptake ($\chi$) on the co-limitation model of autotrophic growth, showing that futile cycling only occurs when both activities are present at high levels. As discussed in the main-text, this quantitative view helped us understand why co-expression of CA and Ci uptake activities was not deleterious to CCMB1 or *C. necator* (Figures 4-5), but rather beneficial to CCMB1, enabling modest growth in ambient air (Figure 6). This understanding relies on a fundamental difference between CA and Ci uptake: that Ci uptake is energized and can work against equilibrium, while CAs are not coupled to any energy source and cannot.

Given that CAs are not energy-coupled, they cannot cause any leakage or futile cycling on their own. This is clearly seen by considering Figure 7C or the bottom row of S14: if CA activity $\delta$ was increased while Ci uptake $\chi$ is kept low, the modeled cell did not leak Ci. At best, CA expression can lead to equilibration of the Ci pools on both sides of the membrane (Figure S14A-B). Based on a variety of experiments, Ci uptake systems are considered to use energy to concentrate $HCO_3^-$ in the cytoplasm either by pumping extracellular $HCO_3^-$ or by energy-coupled hydration of $CO_2$ at the cell membrane. The energy sources used range from ATP to redox and ion gradients (26, 35, 36). Regardless of the underlying mechanism, our current understanding of the CCM requires a high intracellular $HCO_3^-$ concentration that is, crucially, not in equilibrium with $CO_2$ (7, 37, 38). This is understood to be the reason that expression of cytoplasmic CA activity is highly deleterious to photosynthesis and growth in model Cyanobacteria (37).

Energy-coupled Ci uptake can therefore concentrate $HCO_3^-$ in the cytosol and $HCO_3^-$ spontaneously dehydrates to $CO_2$ on a timescale of ≈10 s (7, 22). High $\chi$ values can therefore produce Ci leakage on their own, which can be seen in Figure 7C and S14 where very high $\chi$ values lead to both $CO_2$ and $HCO_3^-$ leakage, i.e. $J_{L,B}$, $J_{L,H} > 0$. Leakage of $CO_2$ indicates that some $HCO_3^-$ dehydrates to $CO_2$, some of which can be used by rubisco. This effect is amplified by CA expression: when $\delta$ was increased at high $\chi$, zero leakage ($J_{L,tot} = J_{L,B} + J_{L,H} = 0$) could be achieved at relatively lower $\chi$ (Figure 7C and bottom row of S14) without altering the flux to biomass ($J_B$) substantially (depicted in log-scale in Figure S14I). According to our model, therefore, modest co-expression of CA and Ci uptake can reduce energy expended on pumping and balance the supply of $CO_2$ and $HCO_3^-$ with the cellular demand for rubisco and bicarboxylation flux.

When $\delta$ and $\chi$ were both set to high values, the model produced substantial futile cycling with $J_{L,tot} / J_B \approx 100$ in extreme cases. First note that these values of $\delta = \chi = 10^3\ s^{-1}$ are several orders higher than the upper bounds we estimated above. Nonetheless, we can ask whether such a leakage rate should be expected to be deleterious to growth by comparing the energy expended on Ci pumping and CO2 fixation. Ci pumping consumes ≈ 1 ATP/carbon (7, 36) while $CO_2$ fixation in the Calvin-Benson-Bassham cycle consumes 2.3 ATP/carbon (39, 40). Therefore, $J_{L,tot} / J_B \approx 100$ implies that 40-50 times more cellular energy is expended on Ci pumping than on $CO_2$ fixation.

### Supplementary Figures

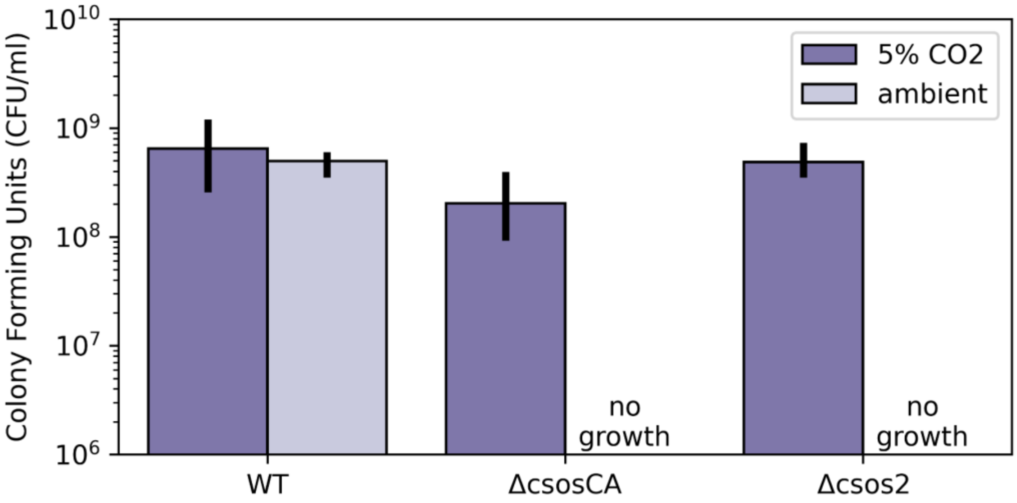


**Figure S1: *H. neapolitanus* CCM mutants grow 5% $CO_2$ but not in ambient air.** Quantification of panel B of Fig. 1. Wild-type *H. neapolitanus* (WT) grows well in 5% $CO_2$ (dark purple) and ambient air (0.04% $CO_2$, lighter purple), producing > $10^8$ colony forming units per milliliter of culture in both conditions. Mutants lacking genes coding for essential CCM components grow in elevated $CO_2$ (dark purple) but fail to grow in ambient air (light purple). The Δ*csosCA* strain lacks the gene coding for the carboxysomal carbonic anhydrase (*csosCA*) while the Δ*csos2* strain lacks the gene coding for an unstructured protein, *csos2*, required for carboxysome formation (41, 42). These mutant strains both failed to grow in ambient air ("no growth"), but grew robustly in 5% $CO_2$ (≈$10^8$ colony forming units/ml). Bar heights give the mean of counts for three biological replicates, which each represent the mean of three technical replicates. Error bars give the standard deviation of the mean. See Table S4 for full description of strains and mutations.

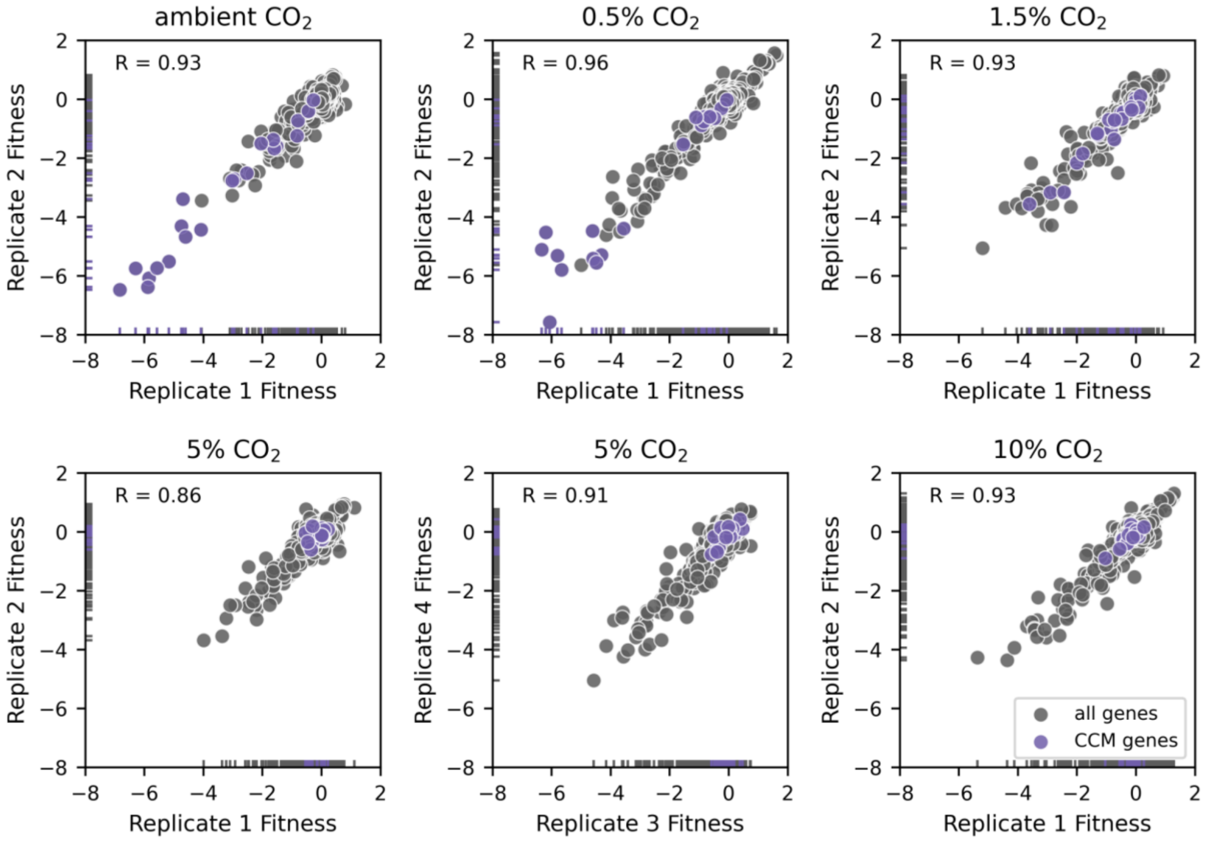


**Figure S2: Reproducibility of *H. neapolitanus* fitness measurements across replicate experiments in the same $CO_2$ environment.** All $CO_2$ conditions were assayed via duplicate cultures with biologically independent pre-cultures, except for the 5% $CO_2$ condition which was assayed in biological quadruplicate. Scatterplots show the correlation between replicates for those genes which produced high confidence fitness measurements in both replicates, with known CCM genes in purple and all other genes in grey. The Pearson correlation R is given for all pairs of replicates plotted and exceeds 0.85 in all cases. Marginal distributions of per-replicate fitness effects are given by the "rug" along the axes. As CCM gene disruptions (purple) represent the largest fitness effects observed in lower $CO_2$ conditions, the range of fitness effects decreases with increasing $CO_2$.

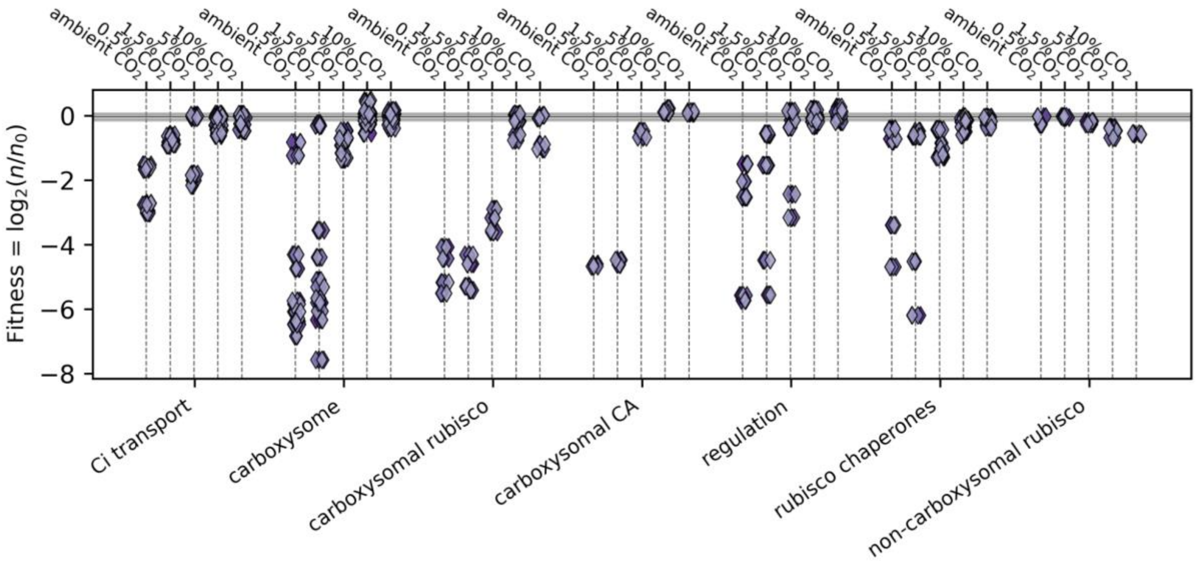


**Figure S3: Contributions *H. neapolitanus* CCM genes to organismal fitness across five environmental $CO_2$ concentrations.** As in Figure 2, data derive from batch competition assays of a barcoded whole-genome insertional mutagenesis library (RB-TnSeq) developed in (26). Data for ambient and 5% $CO_2$ conditions are reproduced from that reference, while data 0.5%, 1.5% and 10% $CO_2$ conditions were collected for this study. Each competition assay was performed in duplicate, except for the 5% $CO_2$ condition, which was performed in quadruplicate (i.e. biological duplicate in each study). We manually divided CCM-associated genes into several categories based on their known or presumed roles. The correspondence between genes and categories is given Table S1. The figure plots the fitness effects of knockouts for each gene category as a function of the $CO_2$ level and include three additional categories of genes omitted from Figure 2: putative transcriptional regulators of the CCM, rubisco chaperones, and the non-carboxysomal Form II rubisco ("non-carboxysomal rubisco"). The presence of a non-carboxysomal rubisco explains why mutations disrupting the carboxysomal enzyme are not very deleterious in 5-10% $CO_2$: the secondary rubisco is expressed in those conditions (43). The interpretation of fitness results is complicated by genetic redundancy for several other gene categories as well. For example, the *H. neapolitanus* genome encodes 6 carboxysome shell proteins, which differ in their abundances (44) and could have overlapping roles in the carboxysome structure (36, 45). Five of these proteins are encoded by genes in the major carboxysome operon (26, 36), which can cause polar effects where the knockout of an upstream gene has a larger effect due to perturbation of transcription of genes encoded downstream (46). Likewise, *H. neapolitanus* has two DAB-type Ci uptake complexes. These complexes are encoded by 2-3 genes each and are both functional when expressed in E. coli (26, 47), which may explain the complex $CO_2$-dependent phenotypes observed for "Ci transport" genes. The "regulation" and "rubisco chaperones" categories are more ad-hoc, as they group multiple genes with poorly-documented roles. Knockout of the rubisco chaperone acRAF, for example, is associated with sizable $CO_2$-dependent fitness defect, though it is as-yet unclear what role this gene plays in rubisco or carboxysome biogenesis in bacteria (1, 48).

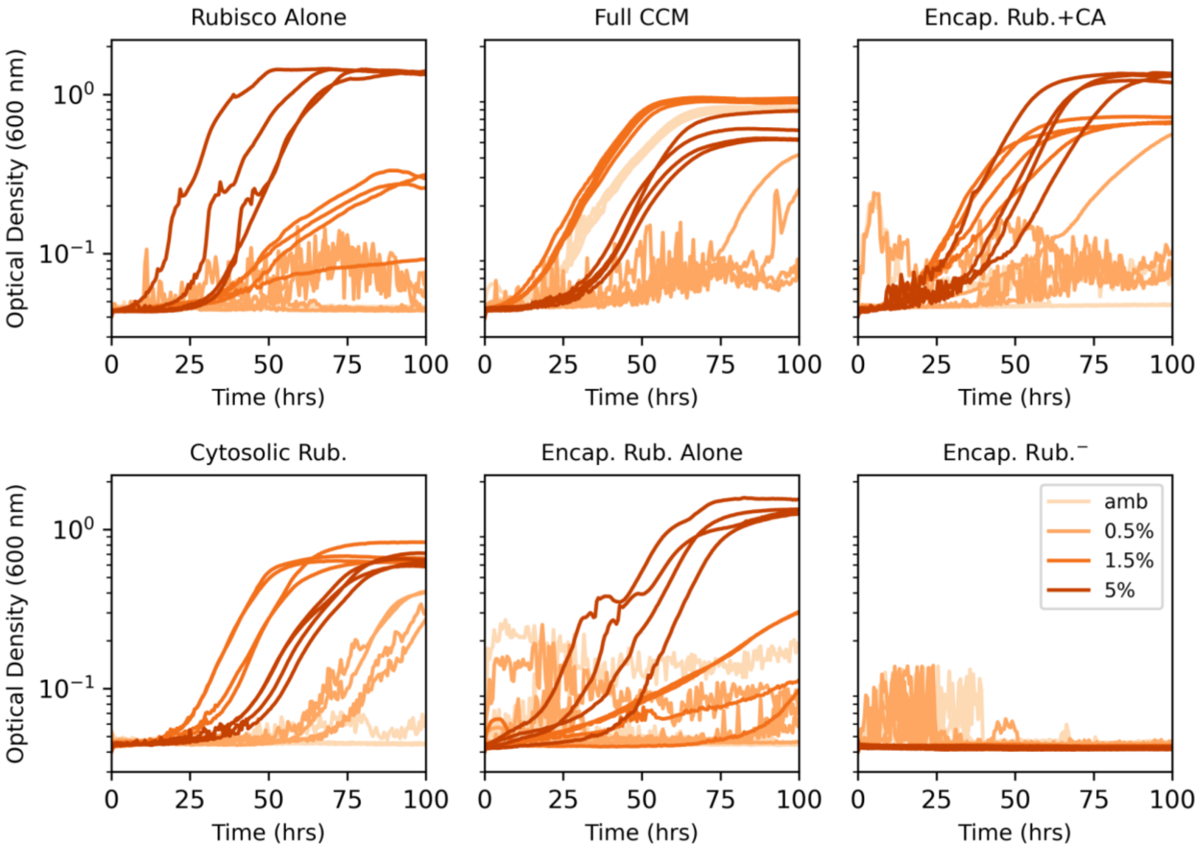


**Figure S4: Growth curves testing the effect of rubisco encapsulation on the growth of CCMB1 in various $CO_2$ pressures.** Each panel displays four biological replicate growth curves grown in four $CO_2$ pressures marked. The $CO_2$ pressure is denoted by the shade of orange in each panel. Figure 3 plots the endpoint densities of these curves (density at 100 hours). The CCMB1 *E. coli* strain grows in elevated $CO_2$ (1.5 and 5%) when rubisco is expressed ("Rubisco Alone", top left). Expressing the full complement of CCM genes ("Full CCM", top middle) permits growth in all $CO_2$ levels. Omitting the DAB-type Ci transporter from this construct ("Encap. Rub. + CA", top right) nonetheless improves growth above the "Rubisco Alone" baseline in 0.5% and 1.5% $CO_2$. Mutating a single amino acid on rubisco (CbbL Y72R) eliminates carboxysome localization by abolishing Csos2 binding (42). Introducing this mutation to a "Full CCM" construct ("Cytosolic Rub.", bottom left) abolishes growth in atmosphere, as reported in (1), but not in 0.5% $CO_2$ or higher. Therefore, carboxysome localization of rubisco is not required for robust growth in 0.5% $CO_2$. Removing carboxysomal CA activity from the "Encap Rub. + CA" construct by active site mutation (CsosCA C173S) abolishes the growth improvement observed when active CA is present ("Encapsulated Rub. Alone", bottom middle). This result implies that the robust growth observed for "Cystolic Rub." and "Encap Rub.+CA" strains was due to the presence of carbonic anhydrase activity. A negative control strain carrying inactive rubisco ("Encap Rub.⁻", CbbL K194M) fails to grow in any condition, as expected. See Table S4 for strains, Table S5 for plasmids and *Methods* for growth conditions.

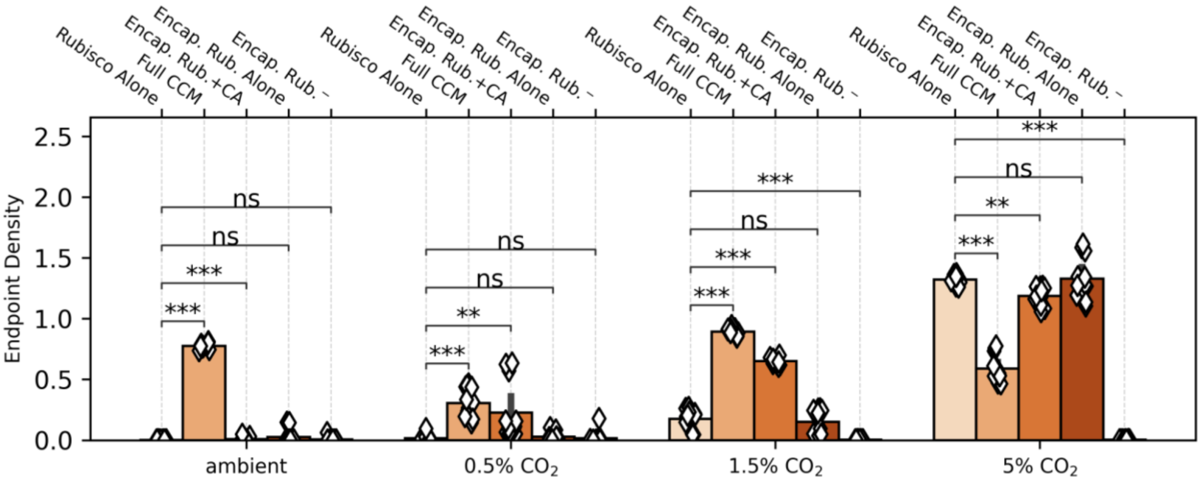


**Figure S5: Assessment of statistical significance of differences in endpoint culture densities for CCMB1 strains testing rubisco encapsulation.** Data and labels are identical to Figure 3, but reordered to group different strains grown in the same $CO_2$ condition. P-values were calculated by comparison to the 'Rubisco Alone' reference strain using a Bonferroni-corrected two-sided Mann-Whitney-Wilcoxon test. '*' denotes $p < 0.05$, '**' denotes $p < 0.01$, and '***' denotes $p < 0.001$. 'ns' denotes 'not significant' at the 5% threshold after Bonferroni correction.

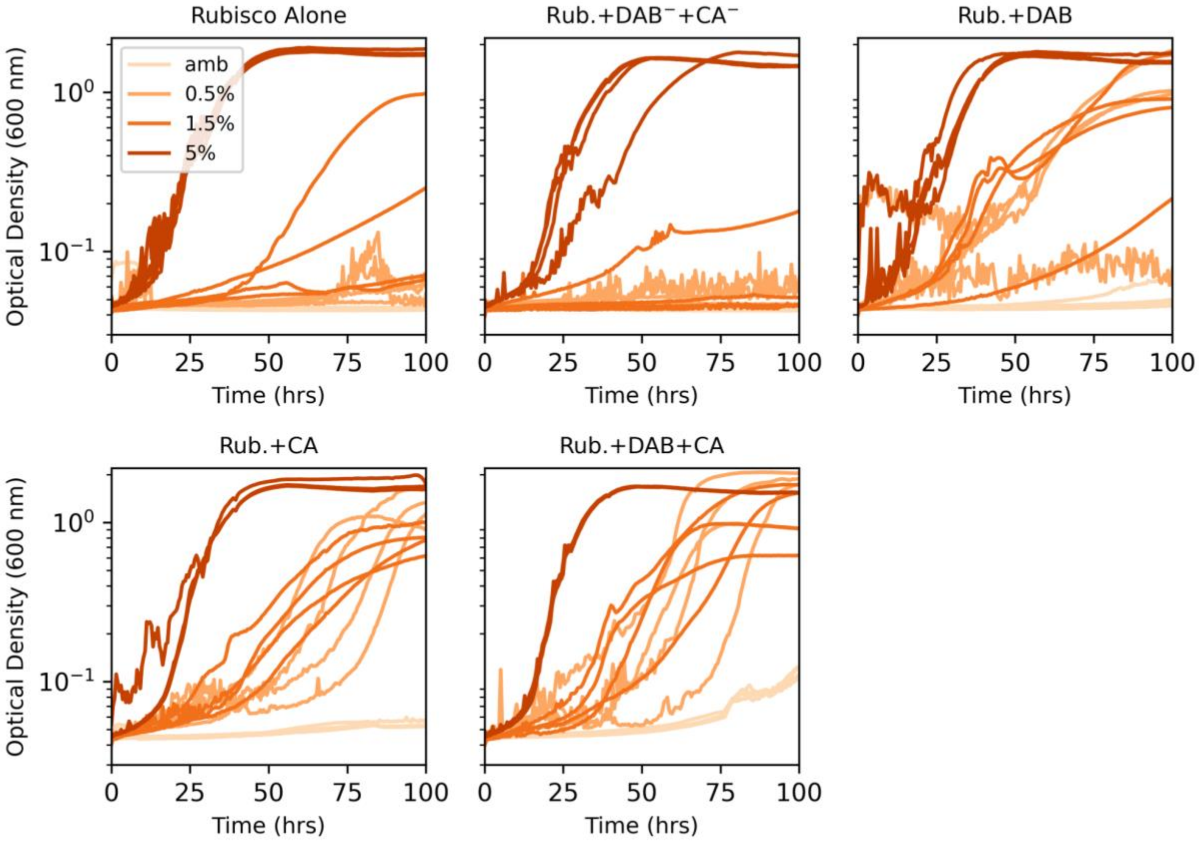


**Figure S6: Growth curves testing the expression of CA and DAB-type Ci transporters on the growth of CCMB1 in various $CO_2$ pressures.** Each panel displays four biological replicate growth curves grown in the four $CO_2$ pressures marked. $pCO_2$ pressure is denoted by the shade of orange in each panel. Labels are identical to Figure 4, which plots the endpoint densities of these curves (i.e. the density at 100 hours).

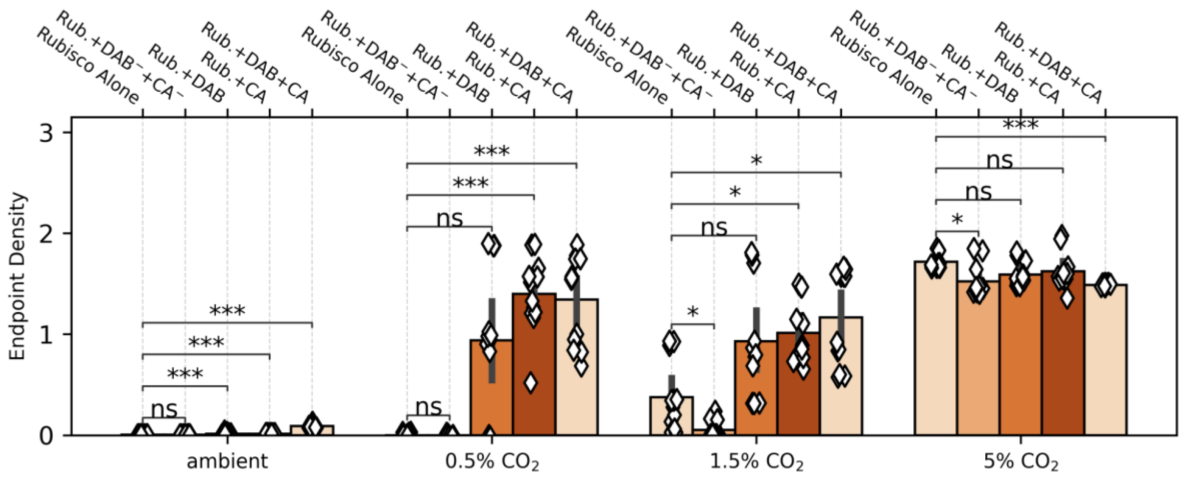


**Figure S7: Assessment of statistical significance of differences in endpoint culture densities for CCMB1 strains testing expression of CA and DAB-type Ci transporters.** Data and labels are identical to Figure 4, but reordered to group different strains grown in the same $CO_2$ condition. P-values were calculated by comparison to the 'Rubisco Alone' reference strain using a Bonferroni-corrected two-sided Mann-Whitney-Wilcoxon test. '*' denotes $p < 0.05$, '**' denotes $p < 0.01$, and '***' denotes $p < 0.001$. 'ns' denotes 'not significant' at the 5% threshold after Bonferroni correction.

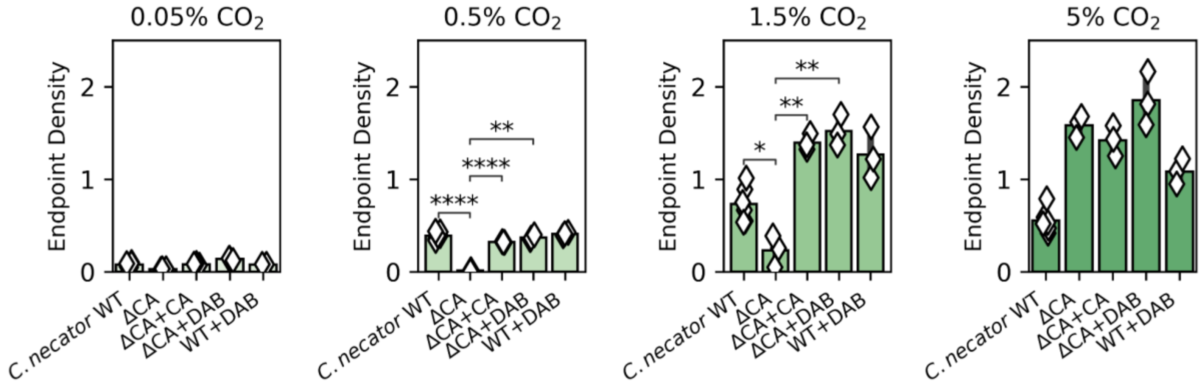


**Figure S8: Assessment of statistical significance of differences in endpoint culture densities for *C. necator* strains testing expression of CA and DAB-type Ci transporters.** Data and labels are identical to Figure 5, but reordered to group different strains grown in the same $CO_2$ condition. P-values were calculated by comparison to the 'Rubisco Alone' reference strain using a Bonferroni-corrected two-sided Mann-Whitney-Wilcoxon test. '*' denotes p < 0.05, '**' denotes p < 0.01, and '***' denotes p <0.001. 'ns' denotes 'not significant' at the P = 0.05 threshold after Bonferroni correction.

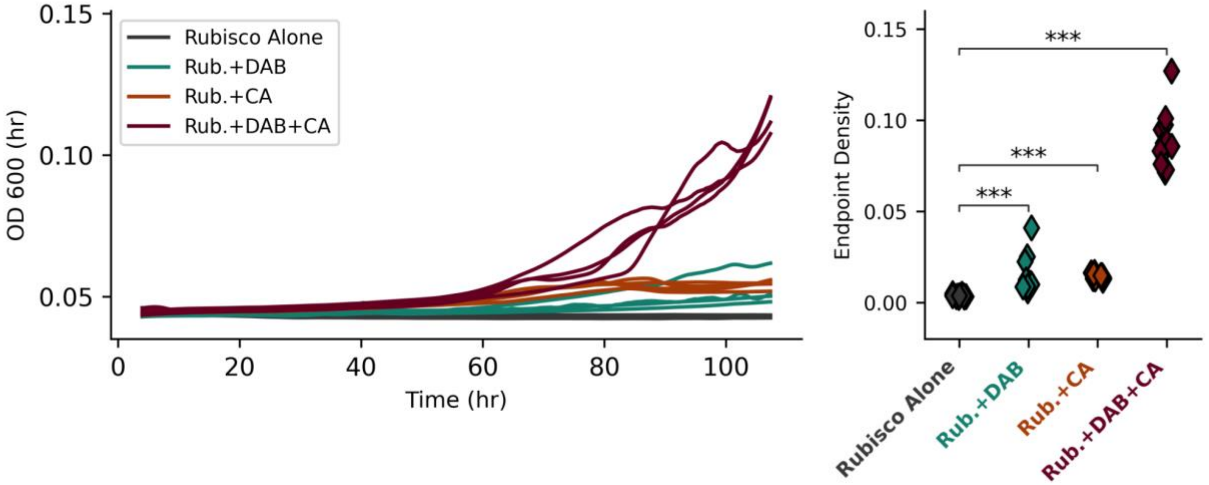


**Figure S9: Growth curves and assessment of statistical significance for CCMB1 strains grown in ambient air.** Left panel gives growth curves for 4 biological replicates of each strain described in Figures 4 and 6. The right panel compares the terminal optical densities for the four strains. P-values were calculated by comparison to the 'Rubisco Alone' reference strain using a Bonferroni-corrected two-sided Mann-Whitney-Wilcoxon test. '*' denotes $p < 0.05$, '**' denotes $p < 0.01$, and '***' denotes $p < 0.001$. 'ns' denotes 'not significant' at the 5% threshold after Bonferroni correction.

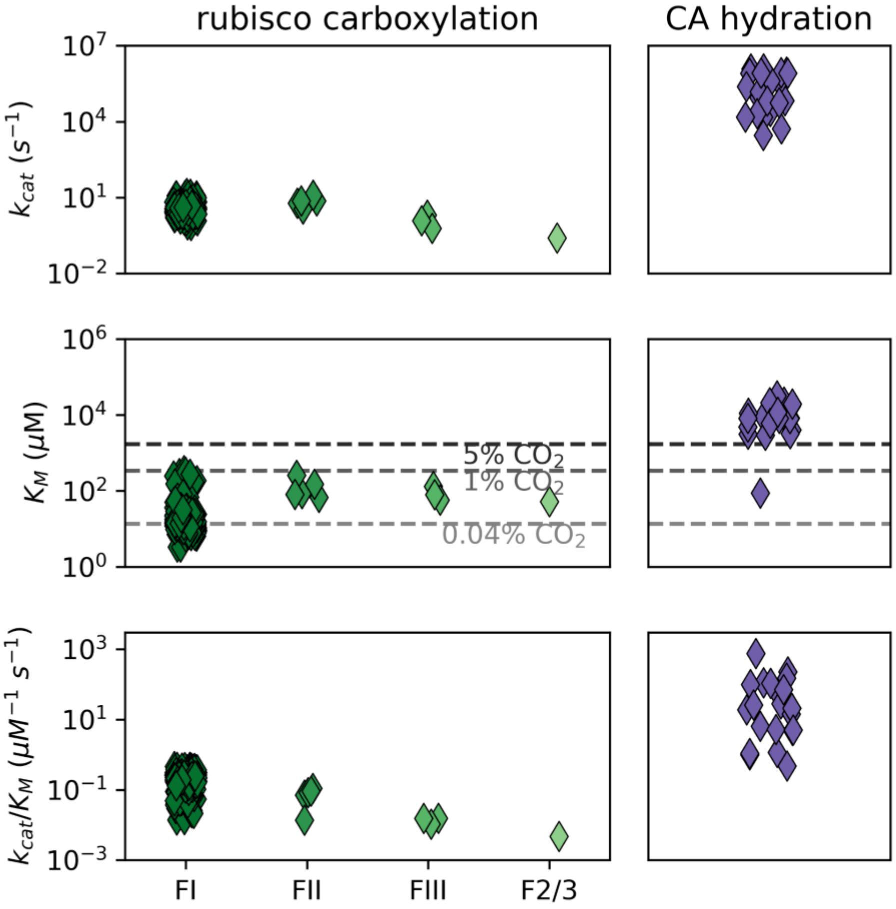


**Figure S10: Literature values of rubisco and carbonic anhydrase kinetic parameters.** In the Michaelis-Menten formalism (49, 50) the $k_{cat}$ gives the substrate-saturated per-active site rate (top panels, $s^{-1}$ units), the $K_M$ denotes the substrate concentration at which an enzyme-catalyzed reaction achieves half the $k_{cat}$ (middle panel, *uM* units) and $k_{cat}/K_M$ gives the per-active site rate in the limit of low substrate concentrations ($[S] \ll K_M$). Rubisco data is drawn from (9) and CA data from (10). Carboxysomal rubiscos are of the form I (FI) variety that is also found in land plants (9, 36). The *H. neapolitanus* genome also encodes auxiliary form II (FII) rubisco. These isoforms typically have higher $k_{cat}$ values, but also lower affinity towards $CO_2$, i.e. higher $CO_2$ $K_M$ values than FI enzymes (51). Less data is available about the kinetics of Form III (FIII) and form II/III (F2/3) rubiscos (52). Notice that $K_M$ values for FI rubiscos are comparable to $CO_2$ concentrations in water equilibrated with present day atmosphere at 25 °C, indicated by the dashed gray line marked 0.04% $CO_2$ (23). Similarly, $K_M$ values associated with CA-catalyzed hydration of $CO_2$ greatly exceed the equilibrium CO2 concentrations. Less data is available about the kinetics of Form III (FIII) and form II/III (F2/3) rubiscos (52). The empirical median $k_{cat}/K_M$ value is 0.2 $uM^{-1}$ $s^{-1}$ (interquartile range 0.17-0.27 $uM^{-1}$ $s^{-1}$) for FI rubiscos and 20 $uM^{-1}$ $s^{-1}$ for CA catalyzed hydration of $CO_2$ (interquartile range 5-98 $uM^{-1}$ $s^{-1}$).

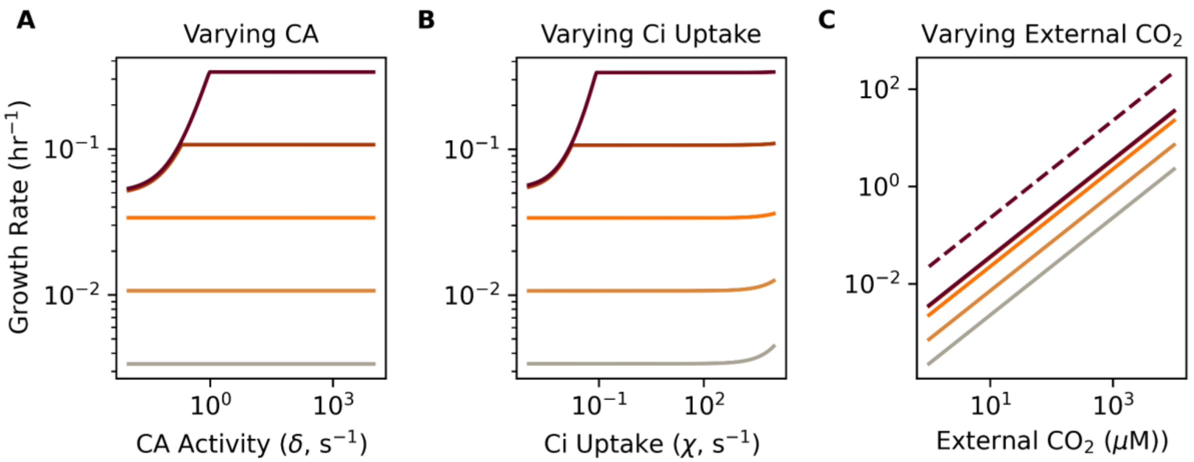


**Figure S11: The effects of individually varying CA activity ($\delta$), Ci uptake ($\chi$), and the extracellular $CO_2$ concentration ($C_{out}$) on growth in the co-limitation model of autotrophic growth.** Panel (A) is identical to main text Figure 7B showing that the model exhibits two regimes: one wherein growth is limited by rubisco flux and another where it was limited by bicarboxylation flux. At low rubisco levels (lighter-colored lines), growth is rubisco-limited: increasing rubisco activity (darker lines) produced faster growth, but the growth rate was insensitive to increasing $\delta$ because slow $CO_2$ hydration provided sufficient $HCO_3^-$ to keep pace with rubisco. At higher rubisco levels (maroon lines), growth was bicarboxylation-limited and increasing $\delta$ was required for increasing rubisco activity to translate into faster growth. (B) Varying Ci uptake activity $\chi$ led to similar effects. As we assume a spontaneous level of $CO_2$ hydration even in the absence of CA ($\delta$ = $10^{-2}$ $s^{-1}$), very high $\chi$ values can increase growth by producing $CO_2$ for rubisco in the rubisco-limited regime. This phenomenon is only apparent at when $\chi$ is implausibly large and the rubisco activity $\gamma$ is small, but is nonetheless instructive for understanding the distinctions between CA and energized Ci uptake. (C) As our co-limitation model is linear, varying the external $CO_2$ concentration produces a proportional increase in the rubisco flux. Additionally, because we assume extracellular $HCO_3^-$ and CO2 are in equilibrium with respect to the pH, $H_{out}$ increases proportionally with $C_{out}$ and supplies sufficient $HCO_3^-$ by passive diffusion and spontaneous hydration of $CO_2$. However, notice that growth does not increase in proportion with rubisco activity as in panels A-B (solid lines represent $\gamma$ values evenly-spaced on a log scale) because, at higher $\gamma = q\omega$ values, passive diffusion and spontaneous hydration of $CO_2$ are insufficient to supply $HCO_3^-$ required for a proportional increase. This can be seen by considering the difference between the solid maroon line (CA $\delta$ = $10^{-2}$ $s^{-1}$) and the dashed one ($\delta$ = 10 $s^{-1}$).

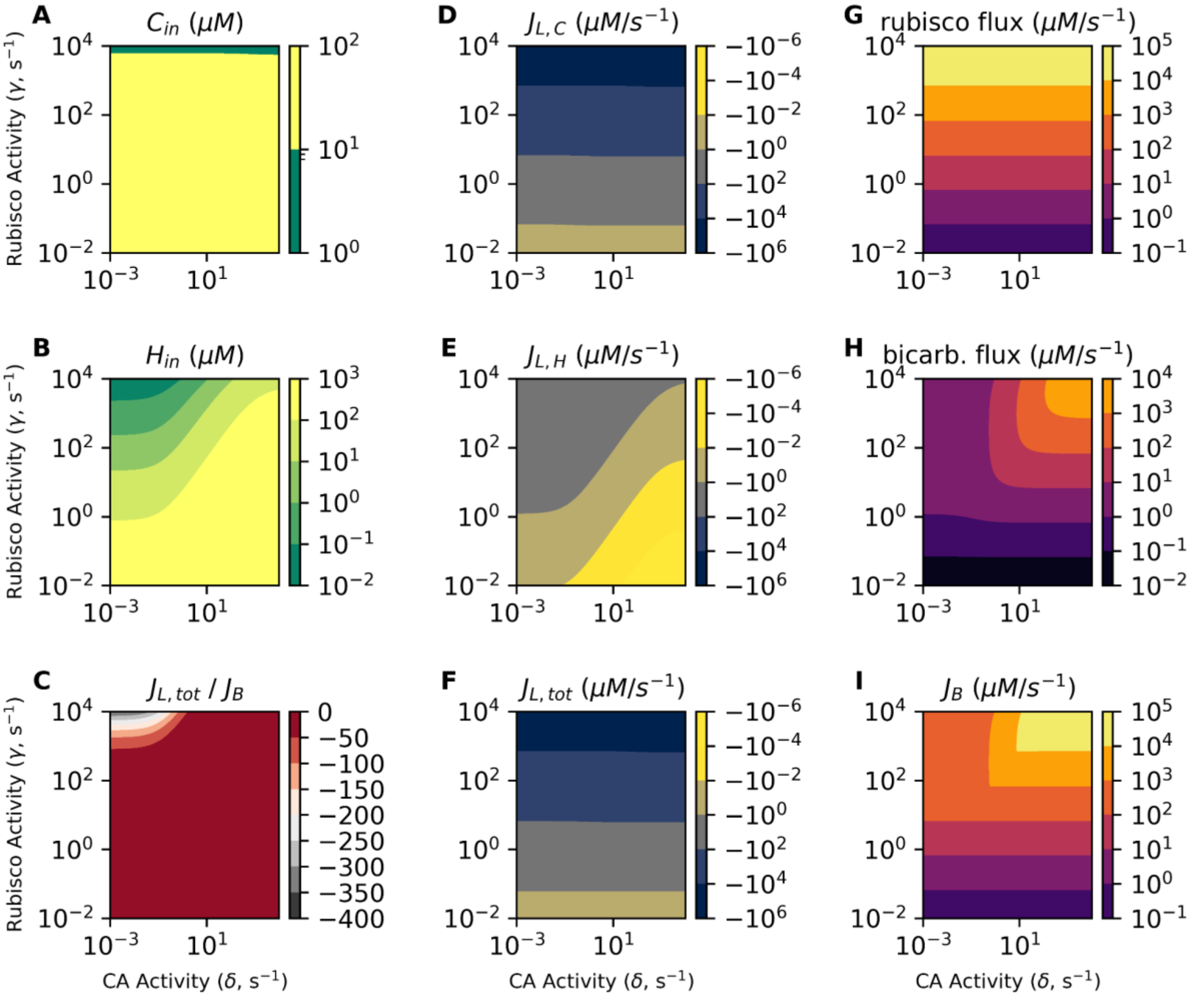


**Figure S12: Rubisco and bicarboxylation-limited growth regimes in the co-limitation model.** In each panel, the x-axis gives the CA activity $\delta$ in s$^{-1}$ units and the y-axis the rubisco activity $\gamma$ in the same units. Color in the filled contour plots gives the quantity named in each panel title. We set $CO_2$ permeability $\alpha$ = 1.2x10$^4$ s$^{-1}$ and $HCO_3^-$ permeability $\beta$ = 1.5 x 10$^{-2}$ s$^{-1}$ as calculated in the supplementary text. The Ci uptake activity $\chi$ was set to 0 for all panels. (A-B) $C_{in}$ and $H_{in}$ are the intracellular $CO_2$ and $HCO_3^-$ concentrations, respectively. Notice that Cin varies little over orders of magnitude changes in $\gamma$ and is independent of CA activity $\delta$ as discussed in the main text. (D-E) $J_{L,C}$ = -$\alpha$($C_{out}$ - $C_{in}$) and $J_{L,H}$ = -$\beta$($H_{out}$ - $H_{in}$) represent the flux of $CO_2$ and $HCO_3^-$ leakage from the cell. $J_{L,C}$ is positive when $C_{in}$ > $C_{out}$ and negative when $C_{in}$ < $C_{out}$ and there is net passive diffusion of $CO_2$ into the cell. As we set $\chi$ = 0, both leakage fluxes are uniformly negative here, connoting passive uptake of both $CO_2$ and $HCO_3^-$. (F) $J_{L,tot}$ = $J_{L,C}$ + $J_{L,H}$ is the total flux of Ci leakage from the cell. Notice that $J_{L,H}$ contributes negligibly to $J_{L,tot}$ here because no $HCO_3^-$ is pumped when $\chi$ = 0. (G) The rubisco carboxylation flux is calculated as $\gamma$ $C_{in}$. Given these permeability values, the rubisco flux is independent of CA activity ($\delta$, x-axis) because passive diffusion of $CO_2$ across the membrane is sufficient to supply even very high rubisco activities ($\gamma$, y-axis). In contrast, panel (H) gives the bicarboxylation flux $\omega H_{in}$, which varies with both $\delta$ and $\gamma$. The dependence on $\gamma$ is an artifact of our assumption that bicarboxylation capacity $\omega$ is proportional to $\gamma$. The dependence on $\delta$ is due to the value of $\beta$, which is low enough that passive diffusion of $HCO_3^-$ across the cell membrane is insufficient at higher $\omega = \gamma / q$. (I) The flux to biomass is calculated as $J_B$ = min($\gamma C_{in}$, $\omega H_{in}$ / $q$). When rubisco activity $\gamma$ is low, $J_B$ is rubisco-dependent, i.e. depends on $\gamma$ but not on $\delta$. When $\gamma$ is larger, however, $J_B$ can be bicarboxylation-limited, i.e. depend on $\delta$ (via bicarboxylation) but not on $\gamma$. Panel (C) gives $J_{L,tot}$ / $J_B$ as a proxy for the energetic efficiency of growth. Here this value is always negative because $J_{L,tot}$ < 0.

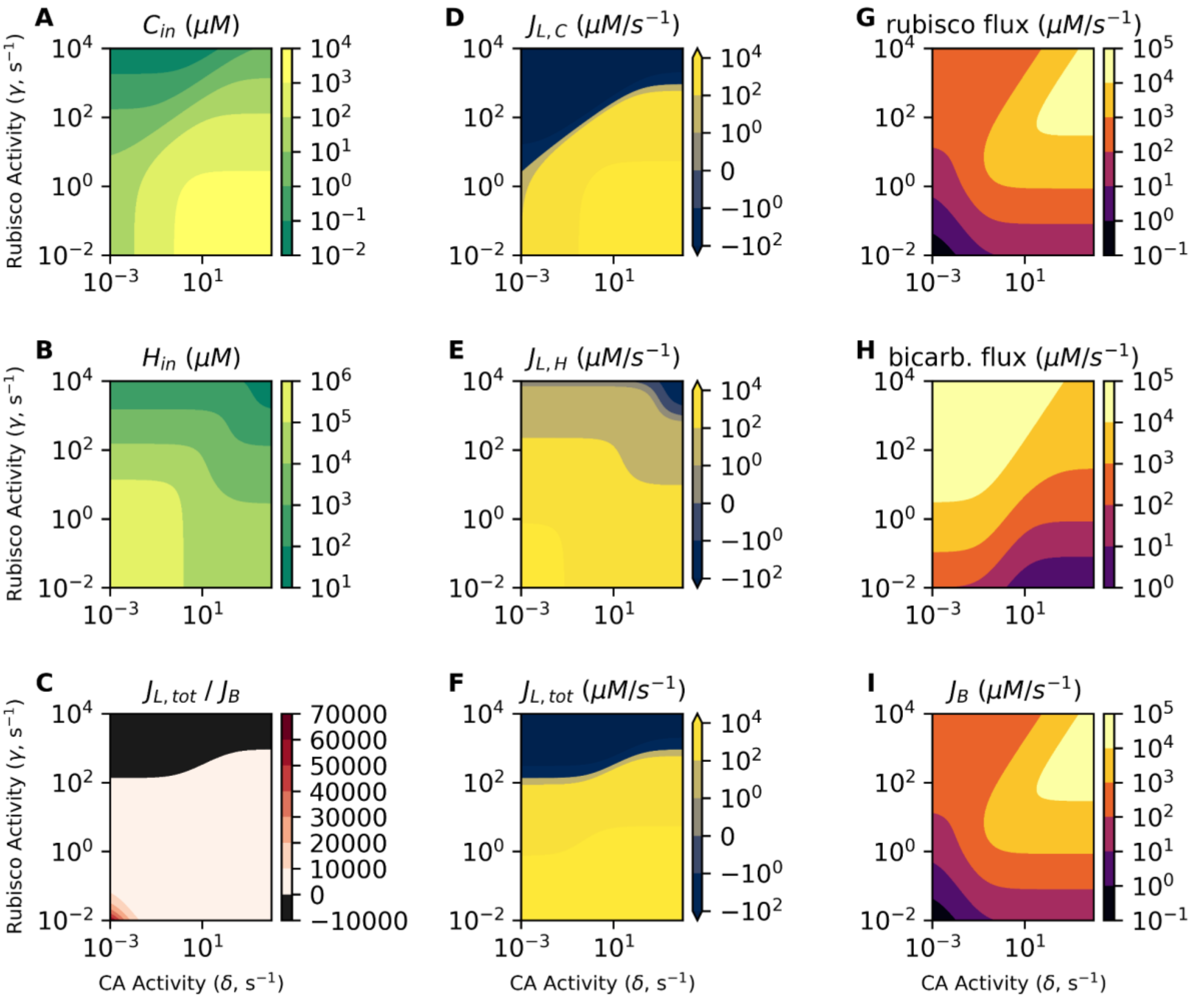


**Figure S13: Unrealistically low $CO_2$ permeabilities permit the co-limitation model to concentrate $CO_2$ intracellularly.** In each panel, the x-axis gives the CA activity $\delta$ [s$^{-1}$] and the y-axis the rubisco activity $\gamma$ [s$^{-1}$]. Color in gives the quantity named in each panel title. Here $CO_2$ permeability $\alpha$ = 12 s$^{-1}$, $HCO_3^-$ permeability $\beta$ = 1.5 x 10$^{-2}$ s$^{-1}$ and Ci uptake activity $\chi$ = 100 s$^{-1}$ for all panels. (A-B) $C_{in}$ and $H_{in}$ give intracellular $CO_2$ and $HCO_3^-$ concentrations, respectively. Given the low $CO_2$ permeability $\alpha$ and Ci uptake capacity $\chi$, it is possible for the model to pump $CO_2$ such that $C_{in} \gg C_{out}$ = 10 uM. (D-E) $J_{L,C} = -\alpha(C_{out} - C_{in})$ and $J_{L,H} = -\beta(H_{out} - H_{in})$ represent the flux of $CO_2$ and $HCO_3^-$ leakage from the cell. As we use a large value of $\chi$, both leakage fluxes can adopt large positive values here. (F) $J_{L,tot} = J_{L,C} + J_{L,H}$ is the total flux of Ci leakage from the cell. Notice that $J_{L,H}$ contributes substantially to $J_{L,tot}$ here because of substantial $HCO_3^-$ pumping ($\chi \gg 0$). (G) The rubisco carboxylation flux is calculated as $\gamma C_{in}$ and depends strongly on $\delta$ because CA activity produces CO2 from pumped $HCO_3^-$ as shown in panel A. Panel (H) gives the bicarboxylation flux $\omega H_{in}$, which also varies with $\delta$ and $\gamma$. The dependence on $\gamma$ is an artifact of our assumption that bicarboxylation capacity $\omega$ is proportional to $\gamma$. The dependence on $\delta$ is due to CA-catalyzed conversion of pumped $HCO_3^-$ (the bicarboxylation substrate) into $CO_2$. (I) The flux to biomass is calculated as $J_B = \min(\gamma C_{in}, \omega H_{in} / q)$. In contrast to Figure S12, biomass flux now depends on $\delta$ even at low rubisco activities $\gamma$. This is due to an unrealistically low value $\alpha$ = 12 s$^{-1}$, which is 1000-fold lower than estimated and measured for biological membranes.

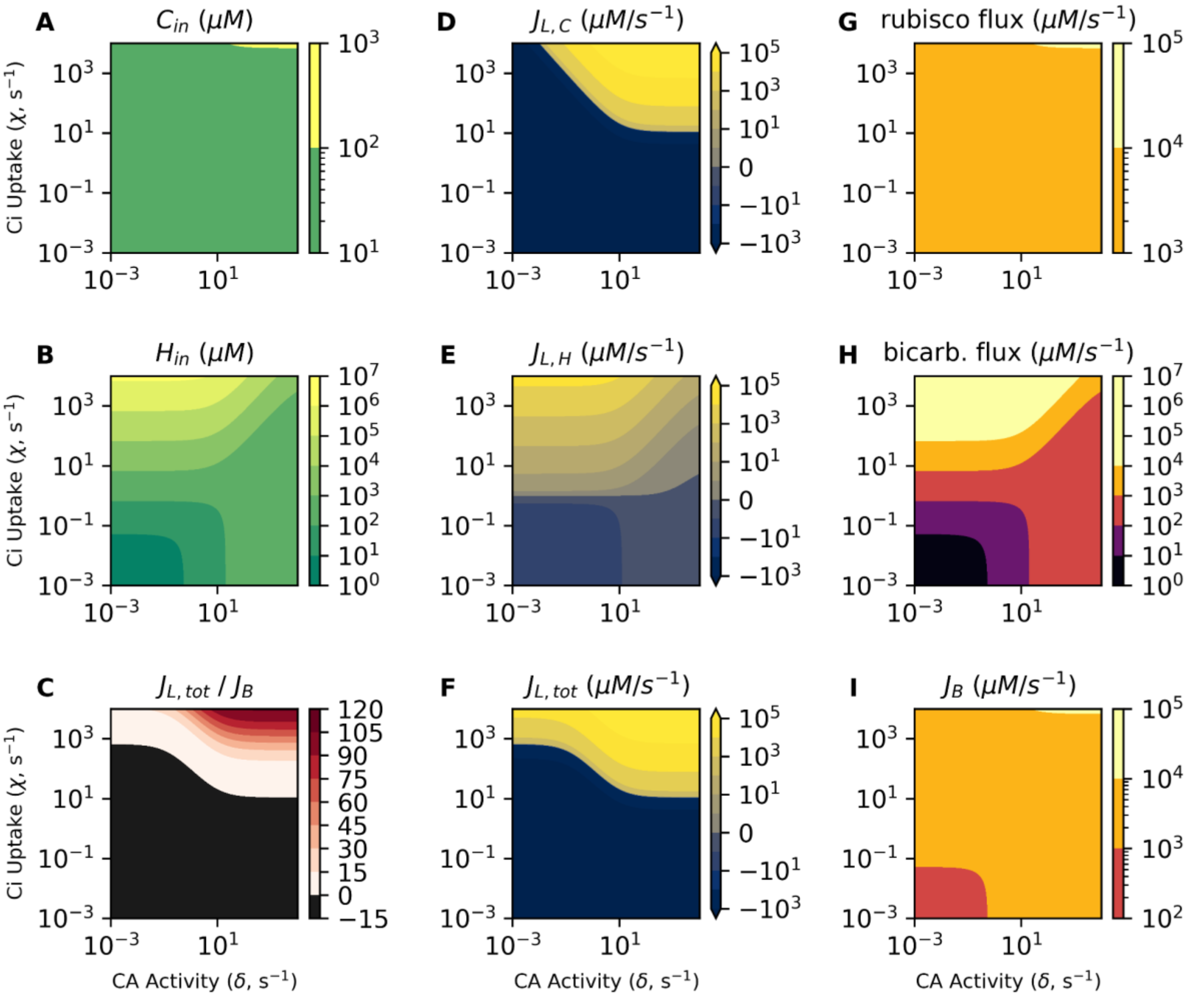


**Figure S14: The effects of simultaneously varying CA activity ($\delta$) and Ci uptake ($\chi$) on the co-limitation model of autotrophic growth.** In each panel, the x-axis gives the CA activity $\delta$ in $s^{-1}$ units and the y-axis the Ci uptake activity $\chi$ in the same units. Color in the filled contour plots gives the quantity named in each panel title. The rubisco activity $\gamma$ was set to 100 $s^{-1}$ for all panels. (A-B) $C_{in}$ and $H_{in}$ are the intracellular $CO_2$ and $HCO_3^-$ concentrations, respectively. (D-E) $J_{L,C} = -\alpha(C_{out} - C_{in})$ and $J_{L,H} = -\beta(H_{out} - H_{in})$ represent the flux of $CO_2$ and $HCO_3^-$ leakage from the cell. $J_{L,C}$ is positive when $C_{in} > C_{out}$ and negative when $C_{in} < C_{out}$ and there is net passive diffusion of $CO_2$ into the cell. (F) $J_{L,tot} = J_{L,C} + J_{L,H}$ is the total flux of Ci leakage from the cell. Notice that $J_{L,H}$ only substantial contributes substantially to $J_{L,tot}$ when $\chi$ is implausibly high; we calculated a maximum value of $\chi \approx 2$ $s^{-1}$ from physiological measurements of cyanobacteria, but values of $\chi \approx 10^3$ $s^{-1}$ are required here for $J_{L,H}$ to contribute noticeably to $J_{L,tot}$ (compare panels D and F). (G) The rubisco carboxylation flux is calculated as $\gamma$ $C_{in}$. Notice that, consistent with our main-text calculation, there is little variation in $C_{in}$ (panel A) and, therefore, rubisco carboxylation (panel G) across orders of magnitude changes in $\delta$ and $\chi$. In contrast, panel (H) gives the bicarboxylation flux $\omega H_{in}$, which varies greatly over the same range due to substantial variation in $H_{in}$ (panel B). (I) The flux to biomass is calculated as $J_B = \min(\gamma\ C_{in},\ \omega H_{in} / q)$. When $\delta$ and $\chi$ are both lo, biomass production is limited by bicarboxylation flux (black region in the lower left) but this limitation is alleviated by increasing either $\delta$ or $\chi$. $J_B$ can be increased further if $\delta$ and $\chi$ are both set to very high values (yellow region on the top right). Panel (C) gives the ratio $J_{L,tot}$ / $J_B$, which is a proxy for the energetic efficiency of autotrophic growth. When $J_{L,tot}$ is large, there is substantial leakage of Ci. This only occurs when $\chi$ is large, meaning that energy is "wasted" pumping Ci that subsequently leaks from the cell. $J_{L,tot} \approx 0$ is desirable because it connotes balance between uptake and carboxylation reactions.
